# Extracellular Vesicles heterogeneity through the lens of multiomics

**DOI:** 10.1101/2024.08.14.607999

**Authors:** Taylon F Silva, Elizabeth Hutchins, Wenyan Zhao, Yari Ciani, Minhyung Kim, Emily Ko, Javier Mariscal, Zhuyu Qiu, Agnes Kittel, Bo Zhou, Yang Wang, Megan Hall, Francesca Galasso, Rebecca Reiman, Michael R Freeman, Sarah Parker, Jennifer Van Eyk, Wei Yang, Edwin Posadas, Jlenia Guarnerio, John Nolan, Clotilde Théry, Andries Zijlstra, Shannon Stott, Sungyong You, Francesca Demichelis, Paul C Boutros, Kendall Van Keuren-Jensen, Dolores Di Vizio

## Abstract

Extracellular vesicles (EVs) are heterogenous in size, biogenesis, cargo and function. Beside small EVs, aggressive tumor cells release a population of particularly large EVs, namely large oncosomes (LO). This study provides the first resource of label-free quantitative proteomics of LO and small EVs obtained from distinct cancer cell types (prostate, breast, and glioma). This dataset was integrated with a SWATH Proteomic assay on LO, rigorously isolated from the plasma of patients with metastatic prostate cancer (PC). Proteins enriched in LO, which were identified also at the RNA level, and found in plasma LO significantly correlated with PC progression. Single EV RNA-Seq of the PC cell-derived LO confirmed some of the main findings from the bulk RNA-Seq, providing first evidence that single cell technologies can be successfully applied to EVs. This multiomics resource of cancer EVs can be leveraged for developing a multi-analyte approach for liquid biopsy.

## Introduction

Extracellular vesicles (EVs) are released by virtually all cell types and play a role in intercellular communication. EVs are heterogenous in biogenesis, size, molecular composition, and biological function^1–4^. EVs originating through exocytosis from the endocytic machinery, historically called exosomes, are generally smaller than 150 nm^5^. EVs originating by direct budding from the plasma membrane (ectosomes), on the other hand, range dramatically in size and can be as small as exosomes or as large as microvesicles^2–5^ and large oncosomes (LO)^6–8^. LO are atypically large EVs (L-EVs) (> 1 μm in diameter) that are released by highly migratory and metastatic cancer cells undergoing amoeboid motility^8,9^. We previously reported that LO isolated from prostate cancer (PC) cells are stronger promoters of growth and tumor angiogenesis than exosomes and achieve this by a MYC-induced reprogramming of cancer-associated fibroblasts^10^. Evidence from an independent team indicates that LO from hepatocellular carcinoma play a crucial role in bone metastasis by promoting an osteoclastic pre-metastatic niche^11^. Another team has reported that LO from glioma stem cells promote disease progression via reprogramming the tumor microenvironment ^12^. Additionally, our group and others have identified cancer-derived LO, which are not detectable in human plasma of healthy controls (*data not shown*), in the circulation of patients with advanced cancer ^2,9,10,13^. Notably, LO have been shown to be particularly abundant in the circulation of glioblastoma patients with short survival time ^12^.

Most EV characterization studies have been performed on small EVs (S-EVs) ^14–17^, the few published studies that have accounted for EV heterogeneity and used large scale profiling to compare EV cargo across cell types have not focused on cancer-derived EVs ^18–20^. Studies integrating proteome and transcriptome of different EV populations are completely lacking, and the scattered reports integrating proteomics and transcriptomics in S-EVs have focused on non-coding RNA using small libraries^19–23^ rather than on mRNA. Additionally, while a massive effort in proteomic profiling of different populations of S-EVs from human and murine samples from different cancer types was recently performed^17^, studies to profile L-EVs across different cancer cell models are completely missing.

We and others have demonstrated that particles in the size range of LO contain more abundant proteins, RNA and DNA cargo when compared to S-EVs ^3,6,13,24^. The fact that LO are associated with disease progression and poor outcome and contain multi-analytes directly derived from cancer cells warrants a deeper investigation. We therefore optimized the isolation of the L-EV fraction that contains LO, focusing on the population of largest EVs, and profiled it comparatively to the S-EV fraction. We performed the first quantitative whole proteome analysis of L- and S-EVs from three cancer cell models to identify EV population specific proteins. We then performed whole transcriptome analysis of PC cell-derived EVs followed by an integrated analysis of protein and transcript levels to identify molecules that are present both at the protein and RNA level and therefore suitable to multianalyte approaches in the future. LO-specific transcripts were then interrogated by single vesicle RNA-Seq, obtained by repurposing a single cell RNA-Seq pipeline to LO. Finally, data-independent acquisition mass spectrometry (SWATH-MS) was performed on LO from PC patient plasma to identify a signature of rapid disease progression, which was used to interrogate the Prostate Cancer Transcriptome Atlas to test correlation with disease progression.

## Results

### LO yield can be readily enriched by low-speed ultracentrifugation

We slightly modified a protocol that was previously applied to human primary dendritic cells but not to cancer cells^15^. The protocol is based on differential ultracentrifugation (dUC) to pellet EVs based on size, followed by floatation on an overlayed iodixanol gradient that allows separation of EVs with different buoyant densities and further purification of EVs from non-vesicular components (**Figure S1A**). Using PC3 cells culture medium, we identified a similar number of large particles in the size range of LO (>1.5 μm) in the low (2,8K) and medium (10K) speed fractions (**Figure S1B**, left panel), but the largest particles (>4 µm size) were significantly more abundant in the low-speed fraction (**Figure S1B**, right panel), suggesting that the largest particles are precipitated at lowest speed. However, the low-speed fraction also contained a few cells (**Figure S1C**), which were eliminated by adding low-speed centrifugation spins (500 x *g*) prior to pellet down the EVs at 2.8K (**Figure S1D-F**). Importantly, these extra steps did not alter the number (**Figure S1G)** or size distribution of EVs as measured with two different technologies (Electrical Sensing Zone (ESZ) and Tunable Resistive Pulse Sensing (TRPS)) (**Figure S1H**). Cell viability prior to EV isolation was close to 100%, indicating that these EVs were not shed by dying cells (**Figure S1J**).

To identify markers of LO and S-EVs across different cancer types, we isolated and characterized three EV fractions from prostate cancer (PC3), glioma (U87), and breast cancer (MDA-MB-231) cells. Even if S-EVs were three orders of magnitude more abundant than L-EVs (**Figure S2A)**, the protein amount was similar in S-EVs and L-EVs (**Figure S2B**), in line with the concept that L-EVs have a larger volume that can accommodate a greater molecular cargo. Both the mean and mode of the size distribution of L-EV were >1 μm and those of S-EV were <130 nm, across the three different cell lines (**Figure S2C**). The presence of LO in the low and medium speed fractions was also confirmed by Transmission Electron Microscopy (TEM) (**Figure S2C**). In summary, we found that a population of L-EVs, comprising LO, can be readily enriched from 3 different tumor cell lines at low centrifugation speed.

### The Large Oncosome signature is evident in the low-speed fraction

Mass spectrometry of the three fractions from the three cancer cell models demonstrated that more than half of the proteins identified in EVs are present in all EV fractions, suggesting most EV protein cargo is shed into all EV types indiscriminately. The number of unique proteins was higher in the fractions collected at low-speed and containing the largest EVs (**Figure S3A)**. The normalized LFQ signal among technical replicates exhibited >95% correlation (**Figure S3B**).

The fraction that contained the largest EVs and the highest number of proteins displayed also the highest total protein abundance (**Figure S3C**). Comparison of ranked normalized protein intensities across the three fractions revealed that EVs collected at medium speed are more likely to have middle-ranking protein intensities than the EVs collected at low-speed EVs and high-speed EVs, which suggests that the middle speed fraction is a mixture of the vesicles present both at low and high-speed fractions. In contrast, both the low and medium speed EVs harbored subsets of proteins with high and low abundances (**Figure S3D**), with similar patterns across the cancer types. Hierarchical clustering revealed a more similar protein profile in the low-speed fraction, which contains LO, across the three cancer models (**Figure 1A**).

**Figure 1.**
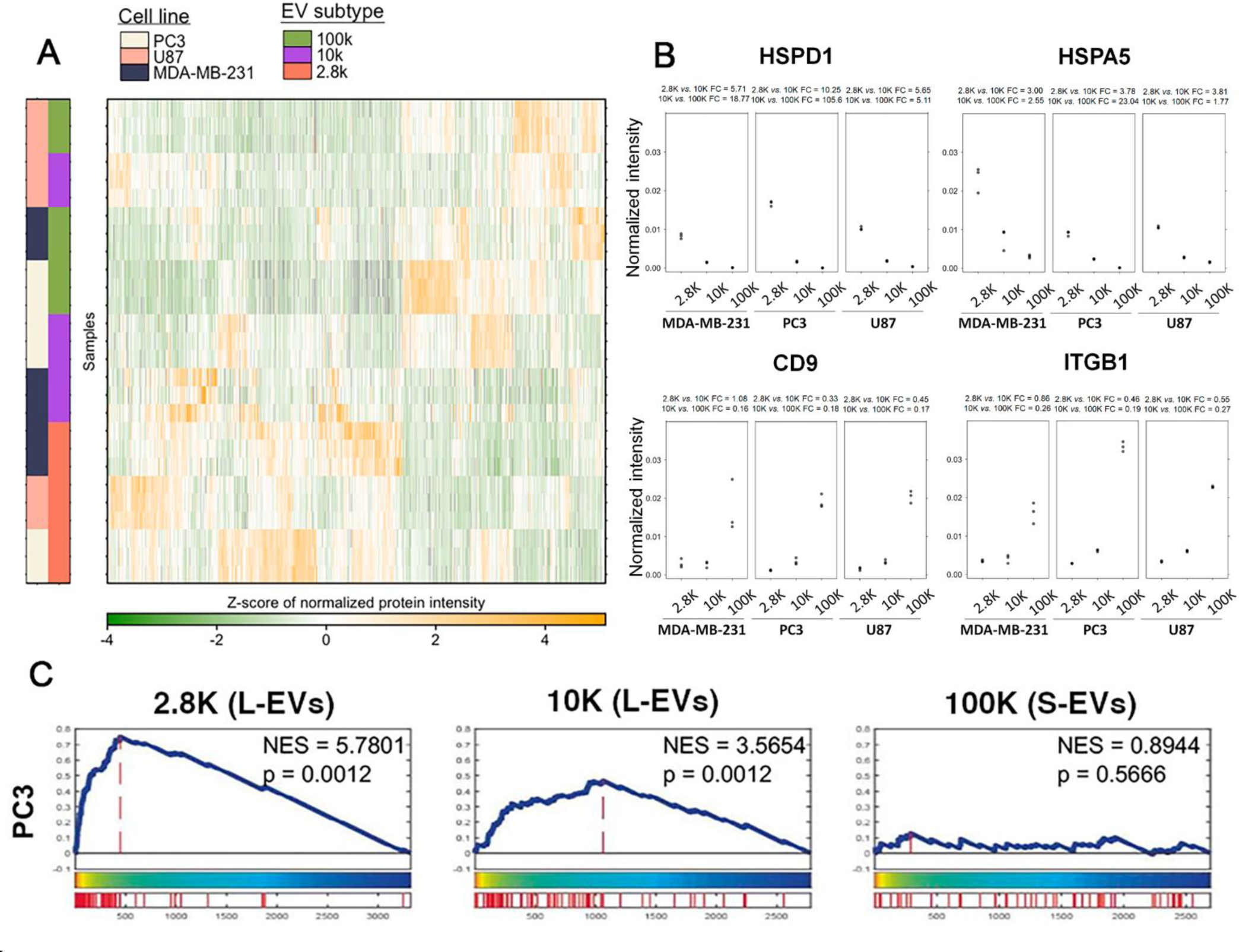
Large oncosome signature is more evident in the low-speed than in the medium speed fraction. (**A**) Divisive hierarchical clustering (DIANA) of the normalized relative abundance of the proteins identified across the indicated EV populations and cell lines. Pearson’s correlation was used to calculate the correlation matrix. (**B**) Normalized relative abundance and fold change of the large EV (HSDP1 and HSPA5) and small EV (CD9 and ITGB1) enriched proteins across the indicated EV pellets and cell lines. (**C**) The gene set enrichment analysis (GSEA) of the proteins differentially enriched in large oncosomes^6^ (FC>1 FDR<0.05) across different EV populations from PC3 cell line. See also Figure S3.

Next, we determined the relative abundance of known LO or exosome proteins in the EV fractions from all cancer models. Proteomics analysis confirmed an enrichment of LO proteins such as HSPD1 and HSPA5 in low-speed versus high-speed fractions across board, highlighting the low-speed fraction as the best source for LO. As expected, S-EV-enriched proteins such as CD9 and ITGB1 were more abundant in high-speed fractions (**Figure 1B**), in line with previous reports^25–27^.

Immunoblotting validated the enrichment of previously identified LO proteins HSPA5 and keratin 18 (KRT18)^6,7,28^ in the low and medium speed fractions compared with the high-speed fraction, confirming that LO can be enriched not only at medium speed, as we and others have previously shown ^2,3,8–10,13^, but even more at low-speed (**Figure S3E**), which has been discarded. In line with rigorous guidelines from the International Society of Extracellular Vesicles, the lack of the intracellular protein Cytochrome C1 (CYC1) in the low-speed fraction, confirmed lack of contamination with cells^29^. Enrichment of TSG101, CD81, and CD9 in the high-speed fraction is consistent with the known enrichment of S-EVs in this fraction (**Figure S3E**). CAV-1 was detected in all EV populations, corroborating our previous finding that this protein, which ubiquitously populates all cell membranes, is present in all EV types^5,28^.

Finally, we looked for an LO signature that was previously identified in medium-speed fractions ^6^ in each EV fraction. Gene Set Enrichment Analysis (GSEA) confirmed an enrichment of the LO signature in medium *vs.* high-speed fractions and revealed an even stronger enrichment of this signature in low-speed fractions, which was particularly evident in the PC cell-derived EVs (**Figure 1C, Figure S3F**). Altogether, our results reveal that a population of LO can be enriched at low centrifugation speed.

### The Large Oncosome fraction from all cancer models is enriched in mitochondrial proteins

Unsupervised clustering identified 104 unique proteins in the low-speed fraction (LO) and 17 unique proteins in the high-speed fraction (S-EVs) across all cancer models. No proteins were identified uniquely in the medium speed fraction (**Figure 2A**). This result suggests that LO and S-EVs can be defined by discrete protein signatures in low and high-speed fractions, respectively. Gene Ontology (GO) analysis demonstrated a dramatic enrichment of mitochondrial components, followed by an enrichment of ER, nuclear, and ribosomal components (**Figure 2B**) in the low speed fraction, which we know is enriched in LO. As expected, the proteins uniquely identified in the high-speed fraction were mostly associated with GO terms such as plasma membrane and extracellular vesicles (**Figure S4A**), confirming enrichment of S-EVs including a population of exosomes in this fraction^14–16^.

**Figure 2.**
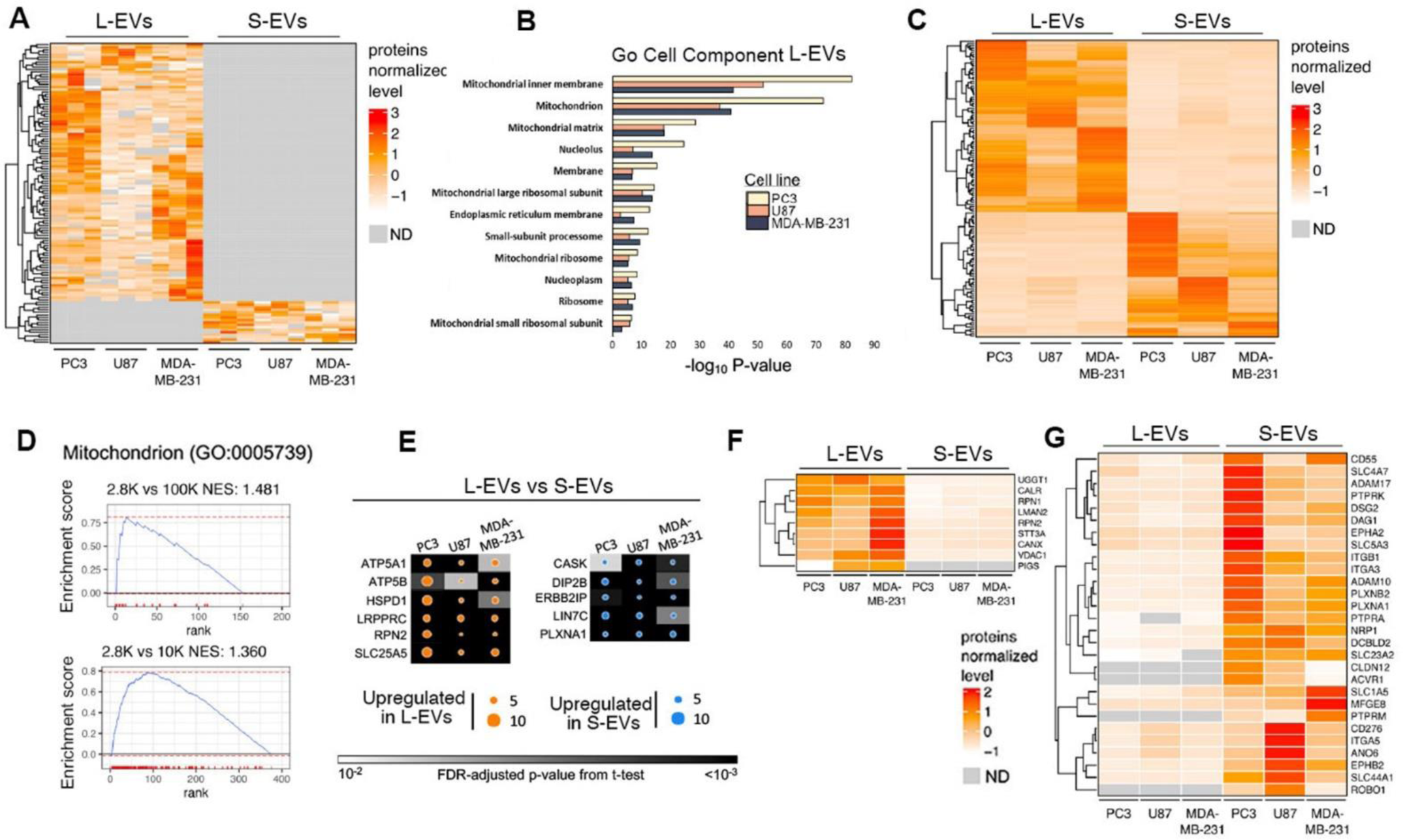
The low and high-speed fractions contain the two most distinct L-EV and S-EV populations. (**A**) Heatmap showing normalized protein intensity of the proteins unique to the low or high-speed EV fractions from the indicated cell lines. ND – not detected. (**B**) Gene ontology (GO) analysis of the proteins unique to the low-speed fraction of all cancer models demonstrates a robust representation of the cell component term mitochondria in L-EVs. (**C**) Heatmap and dendrogram of differentially abundant proteins in L-EVs and S-EVs. ND – not detected. (**D**) GSEA of the differentially abundant proteins using the term Mitochondrion (GO:0005739) showing that mitochondrial proteins are enriched in L-EVs. (**E**) The most enriched (top 25%, fold change > upper quartile) and most highly expressed (top 25%, normalized expression > upper quartile) differentially abundant proteins in L-EVs and S-EVs common to all three cell lines. (**F, G**). Surface proteins unique to or differentially abundant in either L-EVs (**F**) or S-EVs (**G**) and common to the PC3, U87, and MDA-MB-231 cell lines. ND – not detected. See also Figure S3. See also Figure S4.

In line with the unique protein data, the majority of the differentially expressed proteins in LO and S-EVs were more abundant in LO, across cancer types (**Figure 2C, Figure S4B**). GSEA on differentially abundant proteins showed high enrichment for the GO cell component term Mitochondrion (GO:0005739) in the largest EV fraction, in comparison with the other two, further indicating that mitochondrial proteins, which can be found in all EVs, are enriched in LO (**Figure 2D).**

To identify general markers of cancer-derived LO and S-EVs, we focused on the proteins that were not only differentially abundant but also present at high absolute levels in the low and high-speed fractions respectively. The top 25% proteins (fold change and normalized expression) in either the low or the high-speed fraction, common to all three cancer models, were nominated as *bona fide* markers of cancer-derived LO and S-EVs. The majority of the proteins enriched in LO were of mitochondrial origin (ATP5A1, ATP5B, HSPD1, LRPPRC, SLC25A5) with exception of RPN2, which is an ER protein (**Figure 2E, Figure S4C**). Conversely, cell membrane proteins DIP2B, ERBB2IP, LIN7C, and PLXNA1 and the secreted protein CASK emerged as *bona-fide* S-EV markers. The proteins identified in the S-EV fraction have been previously identified in S-EVs both at the protein and mRNA level, deposited in the Vesiclepedia database (data not shown) and can be considered *bone-fide* markers of S-EVs.

Using a published mass spectrometry-derived cell surface protein atlas^30^, we then integrated the current proteomic dataset with the surfaceome and identified several surface proteins in LO and S-EVs from all three cancer models. The number of S-EV surface proteins was higher than the number of LO surface proteins (**Figure 2F, G**). This is consistent with a recent study that, using a mathematical model, reported a higher membrane-to-lumen ratio for S-EVs in comparison to L-EVs^31^. In summary, L-EVs including LO are the EV fraction with more abundant cytosolic and organelle-derived proteins, while S-EVs are enriched in membrane proteins.

### LO markers are informative of response to treatment in metastatic prostate cancer patients

To identify prostate cancer (PC)-specific EV proteins from patients, we attempted to perform quantitative proteomic profiling of LO and S-EVs from plasma samples collected from 20 men with metastatic PC using a highly sensitive, Data Independent Acquisition (DIA) method (SWATH-MS (see **Materials and Methods** for the patient cohort description)). EVs were isolated by differential centrifugation followed by gradient purification. With such a stringent approach, protein in S-EVs was undetectable. Therefore, SWATH-MS could not be performed on this EV fraction. This was not surprising to us, is easily explained by the fact that once the L-EV fraction is isolated, the amount of protein associated with S-EVs is negligible and suggests that what most previous studies have attributed to S-EVs proteins are indeed associated with L-EVs. To investigate whether L-EV markers have prognostic potential, we divided the study cohort into progressive (n = 11) and stable disease (n = 9). For each patient, we analyzed a plasma sample at the beginning of the treatment (T0) and during disease progression or at stable disease (T1) for short or long responders, respectively. To determine whether there were distinct subtypes of EV proteomic profiles across patients, we performed consensus clustering (**Figure 3A**). Three proteomic subtypes, along with four sample subtypes, were identified. The sample subtypes were annotated with clinical features including timing relative to treatment (see **Tables 1** and **2**), PSA levels, prior radiation, and the exposure to other therapeutic strategies. Sample subtypes did not appear to be associated with any specific clinical feature, except for a trend in samples in subtype #3, which had all been exposed to docetaxel. Time-to-event analysis identified 48 proteins associated with time to progression (q < 0.20, Cox proportional hazards modeling) after adjustment for prior ADT and docetaxel. Ten of these proteins (PROS1, NCSTN, CCT2, GNB1, PDZD8, ERLEC1, ARMCX3, C1QBP, RAB38, and RDH14) were associated with shorter time to progression (hazard ratio > 1). Clustering analysis using t-test (*p* ≤ 0.05) identified proteins that were significantly upregulated in progressive vs stable disease (**Figure 3B-C left**). We also identified LO proteins that were significantly upregulated in patients with hormone-resistant versus hormone-sensitive disease at each time point (**Figure 3B-C right**). These findings suggest that LO can potentially be used as a prognostic tool to predict disease progression and treatment outcomes.

**Figure 3.**
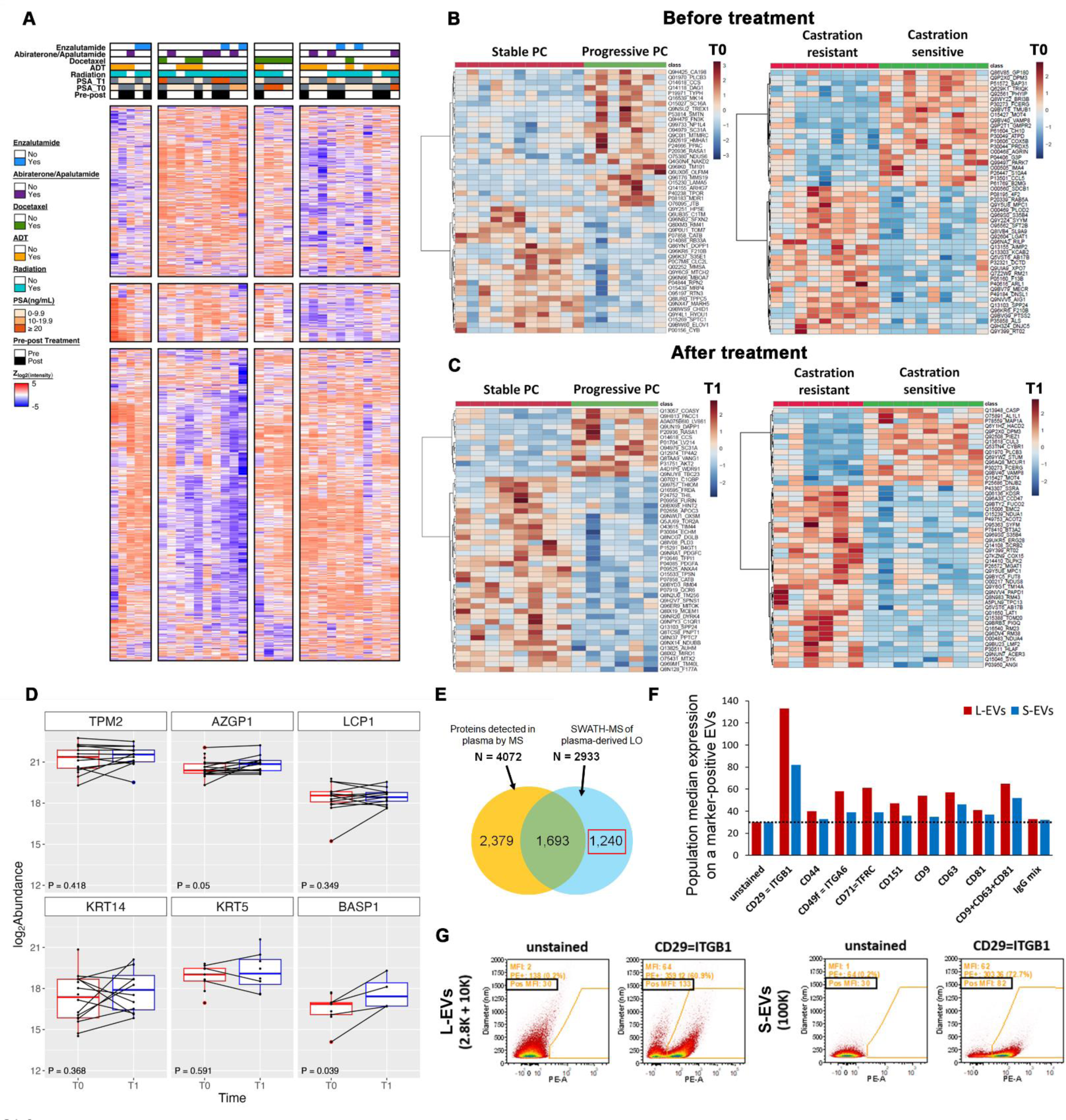
L-EV markers might be prognostic and predictive of treatment outcomes in metastatic prostate cancer patients. **(A)** Consensus clustering of 30 samples (K = 4) using z-scored log^2^(intensity) of the top 75% most variables protein groups that were detected in all samples (K = 3, n = 778). Clinical covariates are shown in the heatmap above, indicating for pre/post treatment, pre-treatment serum PSA Concentration, post-treatment serum PSA Concentration, radiation therapy, androgen deprivation therapy, docetaxel, abiraterone/apalutamide, enzalutamide. (**B-C**) Hierarchical clustering using the Euclidean distance and Ward’s linkage of the proteins differentially expressed (p-value < 0.05) in plasma L-EVs from patients with either stable or progressive PC (left) and castration resistant or castration sensitive PC (right) before treatment (T0) (**B**) and after treatment (T1) (**C**). (**D**) Comparison of log^2^(abundance) of prostate-enriched gene proteins^32^ in matched T0 and T1 samples from plasma patients-derived LO. The plot displays all the proteins detected in our patients-derived LO among these 46 prostate-enriched genes. Box plots show log^2^(abundance) with paired sample differences connected by lines. The P values were calculated using a two-sided paired T test. **(E)** Venn diagram showing the numbers of the L-EV proteins identified in this study that have not been previously detected in human plasma by MS. (**F-G**) Single EV flow cytometry analysis of the select proteins expressed on the EV surface. **(F)** Quantification of the expression of the indicated proteins in molecules per EV. (**G**) Representative dot plots showing the size of the EVs and the expression level of the target proteins.

**Table 1.** Patient characteristics-Castration sensitive prostate cancer (CSPC) cohort. TTP= Time to progression (in months); PSA T0= serum PSA concentration at Time 0; PSA T1= serum PSA Concentration at T1; RT =radiation therapy; ADT = androgen deprivation therapy; LN = lymph nodes; RP = radical prostatectomy

| ID | Treatment | TTP (mo) | Metastatic burden | RP (Y/N) | Radiation | Previous systemic treatment | PSA T0 (ng/mL) | PSA T1 (ng/mL) | T1-T0 (months) |
| --- | --- | --- | --- | --- | --- | --- | --- | --- | --- |
| **CSPC- Treatment Responsive** |  |  |  |  |  |  |  |  |  |
| A1 | ADT | 78.2 | None (prostatic bed recurrence by PSMA) | Y | Salvage RT | none | 0.1 | 0.1 | 4 |
| A2 | ADT + Docetaxel | 60.4 | High (heavy bone + liver) | Y | Salvage RT + metastasis directed RT to bone | none | 0.03 | 0.03 | 5 |
| A3 | ADT + Docetaxel | 36.5 | High (heavy bone + lung) | N | None prior to blood samples | none | 0.4 | 0.61 | 2 |
| A4 | ADT + Abiraterone | 24.2 | High (Bone) | N | Palliative for cord compression | none | 0.04 | 0.06 | 4 |
| A5 | ADT + Abiraterone | 24.8 | High (bone) | N | None | none | 0.14 | 0.27 | 4 |
| **CSPC- Treatment Resistant** |  |  |  |  |  |  |  |  |  |
| B1 | ADT | 8 | Low (LN) | N | None | none | 0.6 | 5.5 | 3 |
| B2 | ADT | 12 | High (LN + bone) | N | None | none | 1.3 | 5.50 | 8 |
| B3 | ADT | 8.3 | High (bone) | N | Primary RT + metastasis directed RT | none | 4.61 | 2.89 | 3 |
| B4 | ADT + Docetaxel | 8.7 | High (bone + LN) | N | Palliative RT to femur | none | 0.45 | 1.96 | 3 |
| B5 | ADT + Apalutamide | 2.8 | High (LN + bone) | N | Primary RT +ADT | none | 25.98 | 384.37 | 3 |

**Table 2.** Patient characteristics-Castration resistant prostate cancer (CRPC) cohort. TTP= Time to progression (in months); PSA T0= serum PSA concentration at Time 0; PSA T1= serum PSA Concentration at T1; RT =radiation therapy; MDT= metastasis directed (radiation) therapy; LN = lymph nodes; RP = radical prostatectomy

| ID | Tx | TTP | Metastatic burden | RP (Y/N) | Radiation? | Previous CRPC treatment | PSA T0 | PSA T1 | T1-T0 (months) |
| --- | --- | --- | --- | --- | --- | --- | --- | --- | --- |
| **CRPC- Treatment Responsive** |  |  |  |  |  |  |  |  |  |
| C1 | Enzalutamide | 86.5 | None (biochemical disease only) | Y | Adjuvant RT | none | 0.06 | 0.07 | 3 |
| C2 | Enzalutamide | 17.8 | Low (LN only) | Y | Adjuvant RT | none | 0.03 | 0.03 | 3 |
| C3 | Abiraterone | 19.5 | Low (bone) | Y | Salvage RT +ADT + MDT | None (progressed during ADT + RT) | 0.03 | 0.03 | 3 |
| C4 | Bicalutamide | 18.1 | Low (bone) | N | Primary RT + salvage HD RT (no ADT) + MDT | Sipuleucel-T, bicalutamide, experimental stem cell therapy (China) | 1.18 | 0.73 | 2 |
| **CRPC- Treatment Resistant** |  |  |  |  |  |  |  |  |  |
| D1 | Abiraterone | 0.9 | High (lung + bone) | Y | Salvage RT (no ADT) MDT | Docetaxel, enzalutamide, MDT (bone), carotuximab + enzalutamide, abiraterone | 16.83 | 29.29 | 1 |
| D2 | Enzalutamide | 0.9 | High (Bone) | Y | Salvage RT | Bicalutamide, apalutamide + abiraterone, MDT (bone), enzalutamide | 16.0 | 31.3 | 1 |
| D3-D | Docetaxel | 0.9 | High (Bone) | N | Primary RT + ADT | Bicalutamide, apalutamide, docetaxel | 71.91 | 96.62 | 1 |
| D3-A | Apalutamide | 7.6 | High (Bone) | N | Primary RT + ADT | Bicalutamide, apalutamide, docetaxel | 0.50 | 13.96 | 6 |
| D4 | Docetaxel | 5.2 | High (bone + LN) | N | none | Enzalutamide, docetaxel | 4.91 | 4.49 | 3 |
| D5 | Docetaxel | 1.8 | High (LN + bone) | N | Primary RT + ADT | none | 774.17 | 781.69 | 3 |

We further performed an analysis using an “*in house*” prostate gene signature composed of 46 prostate-enriched genes identified using RNA-seq data from normal or normal adjacent to tumor (NAT) prostate samples from the Genotype-Tissue Expression project (GTEx) and The Cancer Genome Atlas (TCGA) ^32^. Among the prostate-enriched genes, 6 proteins were enriched in our PC patients-derived LO in both time points analyzed, confirming the presence of prostate-derived proteins in the plasma-derived EVs (**Figure 3D**).

We detected a total of 2,933 proteins in LO samples, over half of which (56.6%) were identified in all patients. The median number of proteins per sample was 2,630±136. Importantly, 1,240 (42.3%) of the LO proteins identified in this study have not been previously detected in human plasma by MS (**Figure 3E**), suggesting that EV enrichment enables identification of cancer cell-derived proteins that are not otherwise quantifiable in unfractionated plasma. This is in line with unpublished data using palmitoyl-proteomic analysis of PC patient plasma, showing palmitoylated proteins in LO from plasma, which could not be detected on straight unfractionated plasma (data not shown).

Using the surface proteins atlas^30^, we assembled a list of candidate surface proteins for PC-derived LO detected not only in PC cell line-derived (PC3) L-EVs but also in patient-derived L-EVs. Some of these proteins were significantly associated with stable or progressive disease) and, therefore, might have a prognostic value on their own: RPN2, PIGS ABCC4, and PLD3 were significantly upregulated in patients with progressive disease and LAMA5, LAMB2, and DAG1 in stable PC patients **(Figure S4B)**. In addition, we identified surface proteins detected in L-EVs from all PC patients, regardless of their response status, and expressed at relatively high levels. Because these proteins were identified in all patients, they can be leveraged for specific isolation of PC-derived LO from patient plasma by immune-affinity approaches. Among the total of 302 surface proteins in the dataset, we identified 39 candidates (**Table S1**) whose relative expression was above the median.

Finally, we used single vesicle flow cytometry to estimate abundance of some of the surface proteins identified in patient-derived EVs on PC3 cell-derived EVs. Using validated antibodies with a calibrated flow cytometer, we found that, as expected, cargo abundance was higher on LO than S-EVs. The median protein abundance was also higher in LO than S-EVs owing to the higher total protein amount per vesicle in LO (**Figure 3F-G**). Notably, protein expression was heterogeneous among EVs, with a fraction of EVs showing measurable antibody staining, while other EVs showed no detectable antigen. The measured single EV antigen expression was generally concordant with MS-based abundance rankings, with a few exceptions (e.g. CD44 and CD9), pointing to the importance of employing orthogonal approaches and understanding the quantitative limits of different methods. In summary, a subset of proteins enriched in LO might be predictive of response to treatment in patients with metastatic PC.

### Whole transcriptome analysis identifies distinct RNA signatures in LO versus S-EVs

We then performed a comprehensive, comparative analysis of the whole transcriptome of LO and S-EVs using the PC model. Normalized counts among the technical replicates for each sample were highly correlated (**Figure S5A**). We found that the length of the RNA transcripts (TPM > 1) with >80% coverage belonging to different biotypes was nearly identical in the PC3 cells and in all three EV populations. However, the frequency of transcripts, which was lower in EVs compared to the cells, was slightly higher in the LO-enriched fraction (**Figure S5B**).

Most of the transcripts (10,074) were detected in all three EV populations and producing cells (**Figure 4A**). In agreement with the proteomic data, each EV fraction had a set of unique transcripts and a set of transcripts shared with other fractions, and unique transcripts were more abundant in the low and high-speed fractions (**Figure 4A**). Principal Component Analysis (PCA) demonstrated that the three EV populations clustered separately from each other and from their producing cells (**Figure 4B**), suggesting that the RNA shed in EVs is not a mere representation of the cellular RNA. Unsupervised clustering analysis of the transcripts differentially enriched in the three EV populations (relative to each other) revealed the most striking differences in the low and high-speed EV fractions, while the medium speed EV fraction was less distinct and appeared transitional between LO and S-EV profiles (**Figure 4C**). This was in line with the proteomic data (**Figure 1A**). Interestingly, both PCA and clustering analysis showed that the mRNA signature of the originating cells was more similar to the low-speed EVs compared to the other two EVs (**Figure 4B, C**), in line with the concept that LO contain several structural cytosolic and organelle-derived molecules that reflect the cell of origin better than S-EVs. In line with proteomics, mitochondrial transcripts were the most enriched (**Figure 4D**) in the low-speed fraction. In summary, whole transcriptome analysis revealed that the low and high-speed fractions contain the most distinct EV populations also at the RNA level, and that LO are enriched in mitochondrial transcripts, in line with the proteomic analysis.

**Figure 4.**
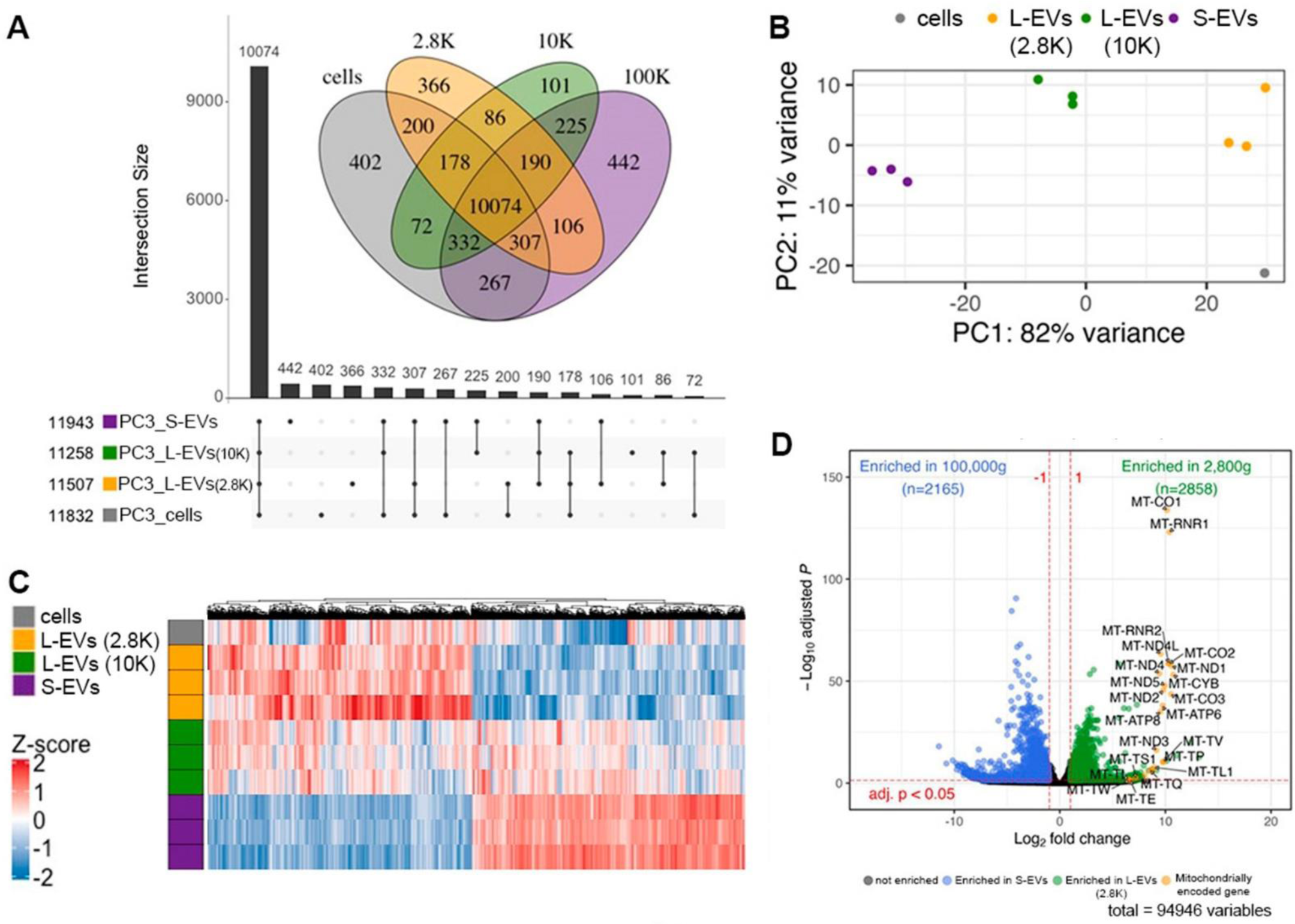
Whole transcriptome analysis identifies distinct RNA signatures in the 2.8K and 100K PC3 EVs. (**A**) Upset plot of the genes identified in the PC3 cell line (DESeq2 normalized counts > 10) between EV fractions and source cells. (**B**) PCA plot of replicates showing grouping by triplicate. (**C**) Heatmap (unsupervised clustering) of Z-scored gene expression of differentially expressed genes between the 2.8K and 100K fractions. (**D**) Volcano plot of differentially expressed genes (adj. p < 0.05, log2 fold change > 1, base Mean > 10) the 2,800g (green) and 100,000g fractions (blue). See also Figure S5.

### Single-EV RNA-seq analysis of LO confirms the abundance of mitochondrial transcripts

To further investigate differential enrichment of mitochondrial transcripts in LO *vs.* S-EVs, we performed GSEA using the GO cell component category for Mitochondrion (GO:0005739). The analysis showed that both low and medium speed fractions were enriched in mitochondrial transcripts in comparison to the high-speed fraction (**Figure 5A**, top and middle). However, a higher enrichment of mitochondrial transcripts was found in the medium speed than in the low-speed fraction (**Figure 5A**, bottom). To determine if these mitochondrial transcripts are measurable in single LO obtained from PC3 cells, we analyzed this fraction using a Chromium 10X technology. To enable encapsulation of single LO into individual droplets, the EV sample was diluted 1:10,000, as typically done for cells.

**Figure 5.**
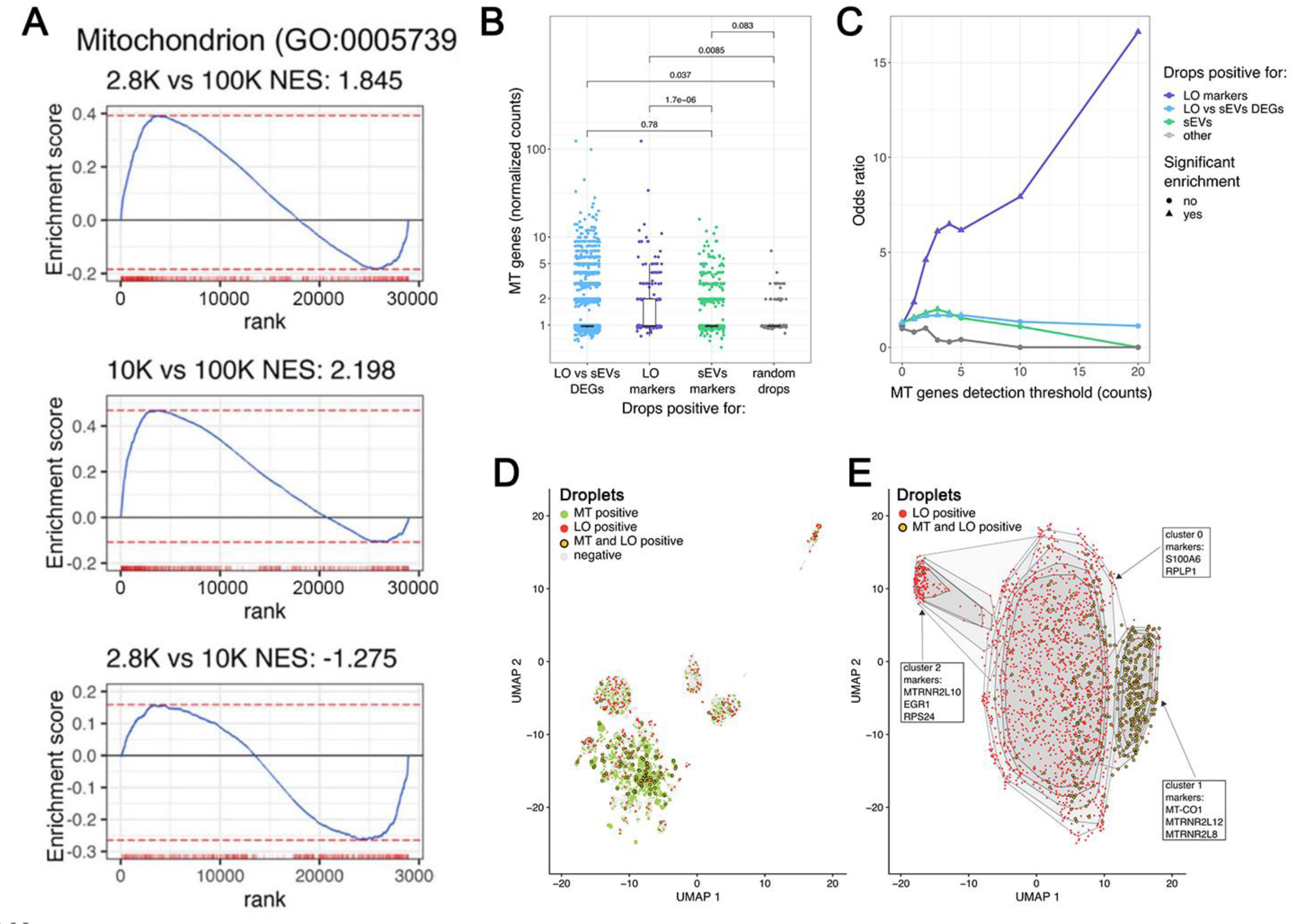
Single-EV RNA-seq analysis of L-EVs confirms the abundance of mitochondrial transcripts. **(A)** GSEA using the GO cell component category for Mitochondrion (GO:0005739) showing an enrichment of mitochondrial genes in both 2.8K and 10K when compared to the 100K (top and middle) and a higher enrichment of mitochondrial genes in the 10K fraction when compared to the 2.8K. Genes are ranked based on Wald’s test statistic of differential expression analysis. **(B)** Expression level of MT genes in droplets positive for: (i) LO vs S-EV DEGs, (ii) LO markers, (iii) S-EV markers, and a set of randomly selected droplets. Statistical significance is obtained with Mann-Whitney two-tailed test. **(C)** Co-occurrence analysis of MT reads and reads of genes classified as (i) LO vs S-EV DEGs, (ii) LO markers, and (iii) S-EVs markers. X-axis represents the number of reads requested to classify a drop as “MT positive”. Y-axis represents the odds ratio of each Fisher’s test, testing the independence of the positivity for MT markers (“MT positive”) and the positivity for other markers (“LO markers”, “LO vs sEVs DEGs”,“sEVs”,“other”). A positive odds ratio indicates an enrichment for EVs positive for MT markers. Significance is shown by the shape of each point, with a threshold of 0.05. **(D)** Dimensionality reduction of single EV RNA sequencing data (only drops containing high number (≥50) of reads are shown). Drops are colored based on the presence of MT and LO transcripts. MT-LO double-positive vesicles (yellow) are observed. **(E)** Dimensionality reduction of single EV RNA sequencing data limited to vesicles positive for LO markers (only vesicles containing high number (≥50) of reads and positive for LO markers are shown). Drops are colored based on the presence of MT and LO transcripts. Clustering analysis identifies 3 clusters. Convex hulls indicate area containing 95%, 90%, 80%, 70%, and 60% of points of each cluster. Significant markers of each cluster are show in text boxes.

Because there are no methods available for the analysis of single-EV sequencing data, we did not use standard data filters but rather applied custom thresholds. Common quality control techniques would, in fact, filter out the signal from single EVs since they contain many fewer transcripts than a single cell (**Table S2**). From the bulk RNA sequencing data, we first obtained signatures associated with LO (LO markers), S-EVs (S-EV markers) and a list of genes differentially expressed between the LO and S-EVs (LO *vs.* S-EV DEGs). We then classified single EVs based on the presence of reads of LO markers, S-EV markers, and LO *vs.* S-EV DEGs (**Figure 5B-D**) and finally investigated the presence of reads associated to mitochondrial transcripts. Globally, 16.3% of droplets contained at least one mitochondrial read (**Table S3**). We observed that the number of normalized counts associated to mitochondrial transcripts was higher in LO than in S-EVs (Mann-Whitney two-tailed test, p = 1.7 x 10^-6^), thus confirming the observed enrichment of mitochondrial signal in LO (**Figure 2D**). To confirm this hypothesis at the single EV level, we performed an enrichment analysis to observe the co-occurrence between LO marker reads and mitochondrial reads. We observed that, independently of the threshold used for the detection of mitochondrial signal (based on the number of reads associated to mitochondrial genes in each EV), droplets positive for LO markers show a significant enrichment for mitochondrial signal (**Figure 5C**). UMAP analysis allowed detection of different clusters (**Figure S6A, Table S4**) of single EVs and the widespread presence of EVs positive for both LO and MT markers (**Figure 5D**). When analyzing only the droplets positive for LO markers we identified a cluster enriched for MT-LO double positive droplets (**Figure 5E and S6B**). Interestingly, the gene MT-CO1 (Mitochondrially Encoded Cytochrome C Oxidase I) was identified as significant marker of the MT-LO enriched cluster (**Table S5**). In summary, LO are suitable for single EV RNA-Seq, which confirmed enrichment of mitochondrial transcripts in LO.

### A protein/mRNA LO signature correlates with cancer progression

Next, we integrated the proteomic and transcriptomic profiles and compared the correlation between the RNA transcripts and their encoded proteins within each EV population as well as between different EV populations. To achieve this, we identified the protein coding RNAs that also had corresponding proteins in our datasets (**Figure 6A**). Most protein coding genes (8,070 out of 11,241) were identified only in the transcriptome and not in the proteome (**Figure S5C**), with only 1,948 genes identified both as mRNA and protein in all three fractions. To determine the EV fraction distribution of transcripts and proteins, we focused on these 1,948 protein coding mRNAs found in both the transcriptome and proteome (**Figure 6A**). The Spearman’s rank correlation between proteins and transcripts in different EV fractions was much higher for transcripts than proteins (**Figure 6B**, blue *vs.* purple squares) and higher in the low (r=0.46) and medium speed (r=0.36) vs the high-speed fraction (r=0.16) (**Figure 6B**, red frames), in line with the idea that EVs with a the larger volume might accommodate both protein and mRNA for the same molecules. Additionally, the mRNAs in the low-speed fraction were slightly more similar to the mRNA in the medium speed (r=0.94) than to the ones in the high-speed fraction (r=0.80) (**Figure 6B**, blue frames). Conversely, the proteins in the medium and high-speed fractions were more similar to each other (r=0.77) than the proteins in low and medium speed fractions (0.66) (**Figure 6B**, purple frames).

**Figure 6.**
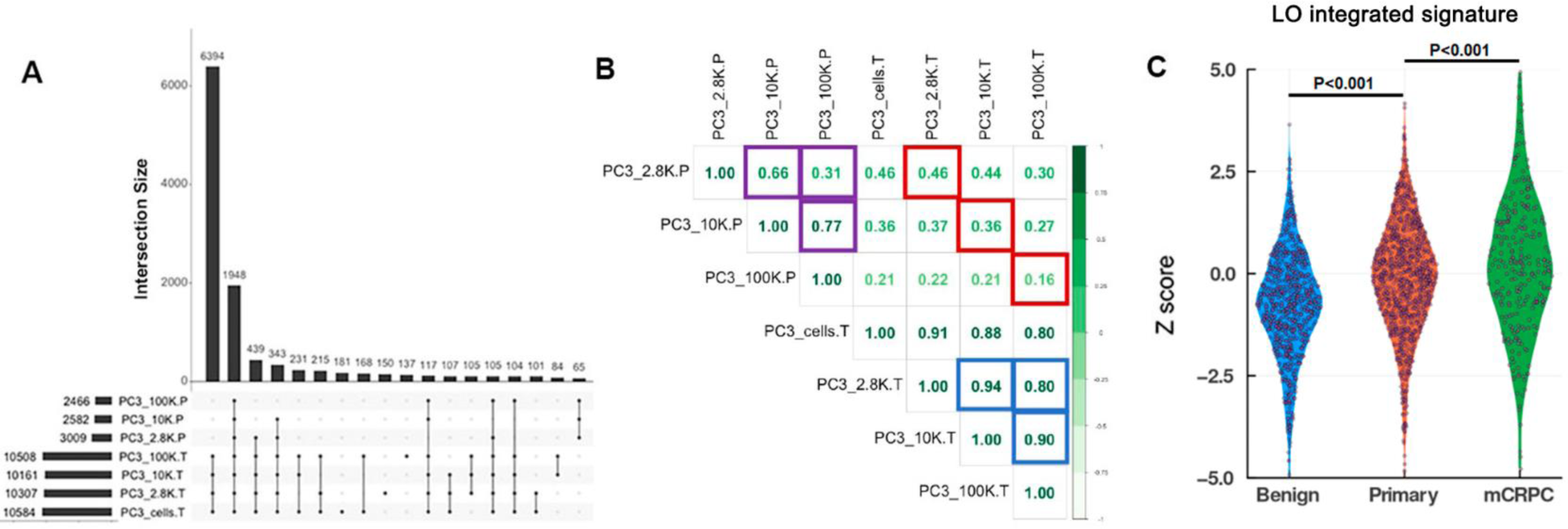
Comparison of protein and mRNA cargo in EV populations in the PC3 cell line. (**A**) Upset plot of protein coding genes identified in the proteome (P) and transcriptome (T; DESeq2 normalized counts > 10). (**B**) Spearman’s rank correlation of genes expressed in both the proteome and the transcriptome. Blue boxes: correlation of different sized EVs within the transcriptome or proteome. Red boxes: correlation of corresponding fractions between the proteome and transcriptome. (**C**) Mitochondrial protein and RNA signature enriched in the PC-derived 2.8K is significantly upregulated in patients with primary cancer *vs.* cancer-free individuals and is further upregulated in patients with mCRPC *vs.* primary cancer. See also Figure S5.

Lastly, we used our integrated proteomic and transcriptomic signatures of the PC-derived LO and S-EVs to interrogate the Prostate Cancer Transcriptome Atlas (PCTA, http://www.thepcta.org/). To generate an integrated signature, we first selected the top candidates (differentially abundant L-EV vs S-EV but also present at high absolute levels) in the EVs from the proteomic analysis (identified both in EVs from cultured cell and from the plasma of patients with metastatic PC) and corresponding protein-encoding RNA candidates from the transcriptomic analysis, resulting in an integrated LO protein-RNA signature: TOMM40, SLC25A5, ATP5B, HSPA9, HSPD1, IMMT, HADHA. These genes were significantly upregulated in primary PC versus benign and in mCRPC versus primary at PCTA analysis (**Figure 6C**). Importantly, all these proteins were highly abundant in LO from the plasma of patients with metastatic PC. The integrated S-EV protein-RNA signature consisted of the following genes: DLG1, GNA13, ERBB2IP, and DIP2B. Expression of these genes did not vary among benign, primary and mCRPC (**Figure S5D**).

## Discussion

Large oncosomes (LO) have been recently acknowledged as key cancer-derived EVs by independent teams ^2,8–12,18^. One of the main advantages is that their atypically large size does not overlap with the size of most of the non-cancer EVs and other non-vesicular particles, including lipoproteins, highlighting a substantial interest in using them for developing EV-based liquid biopsies. This study provides the first resource of large-scale multi-analyte analyses, integrating proteomics and transcriptomics of different populations of highly purified cancer-derived EVs. The results include a source of markers of LO and S-EVs that can be found across three different cancer types. Importantly, these datasets, along with the first single LO RNA-Seq, and the first quantitative SWATH-MS of LO from the plasma of patients with metastatic PC open new avenues for a deeper understanding of EV heterogeneity.

The current literature on LO is scarce and even the literature on S-EVs, which is extensive, often lacks important purification methods to proper separate EVs from other contaminants, like protein aggregates. One of the most commonly used methods for EV isolation is dUC^29^, and usually consist of pelleting L-EVs and S-EVs by medium and high-speed centrifugation, respectively. ^33^Kowal *et al*.^15^ were the first to report the presence of L-EVs at low-speed centrifugation (2,000 x g). Even though these EVs were not comparable in size to LO and were isolated from non-cancer models (dendritic cells), they demonstrated that a low-speed fraction can be a good source for L-EV. Here, using three different cancer models, we demonstrated that the largest EVs, in the size range of LO (1-10 μm), are indeed precipitated with a low-speed centrifugation step and that these particles have a higher LO signature then medium and high-speed fractions, suggesting this as a better method for LO enrichment.

Moreover, LO precipitated in low-speed fractions from different cancer cell models are more similar to each other than different fractions isolated from the same cancer cells, suggesting they might better represent the fraction of circulating cancer material. In addition, those L-EVs are enriched with cytosolic proteins including organelle-derived proteins and metabolic enzymes, with an even higher enrichment for mitochondrial components, both at protein and mRNA levels. Similar to our data, Cocozza *et al*^34^ demonstrated a similar enrichment trend of mitochondrial proteins identified in medium vs high-speed fractions of murine mammary cancer cell line, confirming that mitochondrial proteins are associated with L-EVs.

The enrichment for intracellular organelles, like mitochondria, in LO is corroborated also by RNA-Seq, suggesting that whole organelles are more likely to be in LO than S-EVs. This result might also be a mere reflection of the larger volume of LO, which could randomly incorporate organelles during blebbing of the plasma membrane. Future studies are necessary to investigate potential differences in cancer versus benign cells and whether the organelles are intact or fragmented and if they are functional in recipient cells upon EV transfer. Recent studies reported the presence of mitochondrial material in S-EVs^35–39^; however, none of them compared different EV types or examined L-EVs in the size range of LO. Our demonstration of a significant enrichment of these analytes in LO *vs.* S-EVs is not surprising, given the larger volume of LO vs exosomes. The confirmation of this result by single EV RNA-seq, as a first proof of principle, which showed a significant enrichment of mitochondrial reads only in the droplets containing LO-specific reads, indicates that single EV analysis is possible, at least for this larger population of EVs. Although it is not possible to rule out the presence of more than one LO in each droplet, or the contamination from S-EVs in droplets containing LO, our study opens new avenues for a deeper understanding of the EV heterogeneity.

The finding that LO are enriched in a distinct fraction containing most of the intracellular organelle-derived proteins and transcripts confirms initial reports suggesting that LO are significantly different, at a molecular level, from S-EVs, including exosomes^6^. This is corroborated not only by the findings in the current study that the whole transcriptome profile of LO is more similar than the profile of S-EVs to the originating cells and that the abundance of RNA transcripts with >80% coverage is slightly higher in L-EVs than S-EVs, but it is also in line with a study reporting that while transcripts enriched in EVs are overall shorter than transcripts enriched in cells, larger transcripts are more abundant in L-EVs than S-EVs^40^, which suggests the notion that L-EVs reflect the cellular transcriptome better than S-EVs. However, the integrative analysis of proteome and transcriptome highlights a poor correlation between mRNA and protein in EV fractions. This per se is not surprising, considering the poor correlation between protein and transcript in cells and tissues^41^. However, we show for the first time that this correlation is higher in LO than in S-EVs. This result might be a mere consequence of LO recapitulating cellular profiles better than S-EVs. However, the observation that LO are enriched with ribosomal proteins, and the presence of polyribosomes previously reported in LO^9^ suggest that LO might be an extracellular site for active protein translation, a hypothesis that can be investigated in the future.

On the other hand, the results that more than half of the proteins identified in EVs are present in all EV fractions suggests most EV protein cargo is shed into all EV types indiscriminately but could also be an indication of the limitations of dUC in separating different classes of EVs ^42,43^. Therefore, even though LO emerge as an ideal source of circulating biomarkers to be probed with multi-analyte platforms, better methods for EVs isolation are needed. We have generated a new surface proteins signature from the integrated analysis of highly abundant proteins and protein-coding transcripts detected in LO from different cancer models in LO from mCRPC patient, providing a source for development of immune-capture methods that can be applied to clinical samples and used as diagnostic tool in the future.

In addition, since the set of proteins enriched in LO is common to the three cancer types, they could be utilized as general cancer markers, but this will have to be validated using larger cohorts of patients.^6,44316,4464035–39419^ Our findings in the plasma of cancer patients, showing an association of specific LO cargo with metastatic disease and with a shorter time to progression, highlight the potential value of LO in liquid biopsy. Importantly, our integrated surface signature can be used to interrogate clinical specimens by microfluidics, flow cytometry, immune-capture or other antibody-based technologies that aim to select a population of EVs with a distinct role in cancer. We could also identify a list of proteins in LO associated with a shorter time to progression in our cohort. Additionally, the signature generated from the integrative analysis of the proteins detected in LO of both patients and cancer cell lines with the transcriptome was associated with metastatic disease in the PCTA database.

In summary, we have demonstrated that we can recover a larger number of LO in a specific fraction, enabling their enrichment for a multi-analyte characterization of cancer-derived EVs. Our study provides a source of markers for rigorously collected LO and other EV types that can be utilized for biomarker identification from multi-analyte signatures that might have prognostic value in patients with PC and possibly with other cancer types. More studies are needed to validate these candidates in multiple cohorts of patients and further identify cancer type-specific markers of distinct EV populations.

## Acknowledgements

We would like to acknowledge the mass spectrometry data acquisition and data analysis carried out by the Cedars-Sinai Medical Center Proteomics and Metabolomics Core.

## Author contributions

Conceptualization, T.F.S., D.D.V.; Methodology and Validation, T.F.S., E.H., W.Z., Y.C., E.K., Z.Q., M.K., S.Y., J.M., F.D., P.C.B., K.V.K.J.; Formal Analysis, E.H., W.Z., Y.C., E.K., Z.Q., M.K., S.Y.; Investigation, T.F.S., J.M., A.K., J.N., S.P., M.H., F.G., R.R., B.Z., Y.W., W.Y.; Resources, E.P.; Writing – Original Draft, T.F.S., E.H., W.Z., D.D.V.; Writing – Review & Editing, M.R.F., J.V.E., J.G., C.T., A.Z., S.S., F.D., P.C.B., K.V.K.J.; Visualization, T.F.S., D.D.V.; Supervision, P.C.B., K.V.K.J., D.D.V.; Project Administration, D.D.V.; Funding Acquisition, D.D.V.

## Material and Methods

### Cell culture

All cell lines were obtained from American Type Culture Collection. Cell line identity was validated using STR analysis. Cell lines were maintained at 37°C in 5% CO2. PC3 and MDA-MB-231 cells were grown in Dulbecco’s Modified Eagle Medium (Invitrogen), and U87 cells were grown in Minimum Essential Medium (Invitrogen). The media for all cell lines were supplemented with 10% fetal bovine serum (Denville Scientific), 2 mM L-glutamine (Invitrogen) and 1% PenStrep (Invitrogen). Cell viability of the EV-producing cells was tested with the 0.4% Trypan Blue (Sigma) exclusion method. All cell lines were routinely tested for mycoplasma contamination by using the MycoAlert PLUS Mycoplasma Detection Kit (Lonza). Cell viability was assessed using Trypan Blue Solution, 0.4% (ThermoFisher).

### Enrichment of EVs from cell culture media

Cells were plated into 150 mm tissue culture dishes (n = 18 per cell line) and allowed to proliferate in serum-supplemented media for at least 48 h. After cells reached ∼90% confluency, they were washed twice in PBS, and placed in serum-free media for 24 hours before EV harvest. The enrichment and density gradient purification of EVs was performed as previously reported^6,13,15,28^ with minor modifications described below. The conditioned media was centrifuged three times at 300 *g*, for 5 minutes each to pellet down floating cells. This was followed by centrifugation at 2,800 *g* for 10 minutes to collect the first fraction. The supernatant was spun in an ultracentrifuge at 10,000 *g* for 30 minutes (k-factor 2547.2) to collect the second fraction, and the supernatant was then spun at 100,000 *g* for 60 minutes (k-factor 254.7) to collect the third fraction. All differential centrifugation steps were performed at 4°C. All pellets (2.8K, 10K, and 100K) were then subjected to iodixanol (Optiprep™, Sigma) density gradient purification. Freshly pelleted EVs were resuspended in 0.2 µm-filtered PBS and deposited at the bottom of an ultracentrifuge tube (Cat#344058, Beckman Coulter). Next, 30% (4.3 mL, 1.20 g/mL), 25% (3 mL, 1.15 g/mL), 15% (2.5 mL, 1.10 g/mL), and 5% (6 mL, 1.08 g/mL) iodixanol solutions were sequentially layered at decreasing density to form a discontinuous gradient. Separation was performed by ultracentrifugation at 100,000 *g* for 230 minutes (4°C, k-factor 254.7) and L-EVs derived from the 2.8K and 10K fractions separated into the 1.10–1.15 g/mL density fractions while S-EVs derived from the 100K fraction separated into the 1.10 g/mL density fraction. Purified EVs were then washed in PBS (100,000 *g*, 60 min, 4°C) and resuspended in 0.2 µm-filtered PBS, or in 25 µL of DTT-free 4% SDS/Tris-HCl lysis buffer^44,45^ for proteomic analysis, or in 1 µL TRIzol for transcriptomic analysis. All ultracentrifugation spins were performed in a SW28 swinging rotor (Beckman Coulter).

### Enrichment of large EVs (L-EVs) from patient plasma

All human subject research was performed under Cedars Sinai Medical Center Institutional Review Board-approved protocols Pro00033050 and Pro00042197. These were observational, pilot studies that assess the feasibility of measuring blood-based biomarkers in serially collected blood samples from subjects receiving care within the Cedars-Sinai Center for Uro-Oncology Research Excellence clinics. Subjects provided serial specimens at standard clinical visits with timing set by the treating clinicians. As an observational study, no interventions were mandated by study. Samples were collected only after the participants provided written informed consent. The samples utilized in this study were collected between January 2016 -December 2020. We included two groups of metastatic PC patients: hormone-sensitive (n = 10) and hormone-resistant (n = 10). Samples were selected based upon the availability of plasma before and during therapy to the point of progression mandating a change in treatment. Selected patients were identified as hormone-sensitive if they were treatment naïve and had serum testosterone concentrations > 250 ng/dL prior to therapy then exhibited a response (biochemical and/or clinical) while on hormone therapy. Patients were identified as hormone-resistant if they exhibited biochemical progression with or without radiographic progression while undergoing effective hormone therapy (*i.e.* serum testosterone < 50 ng/dL). Patients were further classified as long or short responders based upon duration of response/benefit from therapy. Details of the subjects are presented in **Tables 1** and **2**. For each individual, plasma aliquots utilized for L-EVs isolation were obtained from the blood draws at 2 separate time points. T0 was selected from a timepoint within 3 months of starting the assigned therapy (noted on **Table 1** or **2**). T1 was acquired three months later, at a point when short responders were already found to progress, and long responders did not exhibit signs of progression. Details of timing are presented in **Tables 1** and **2**. Blood samples were collected in BD Vacutainer™ ACD tubes and processed within 2 hours of phlebotomy. Platelet-poor plasma was prepared from blood by centrifugation at 2000 x *g*, 4°C, for 5 minutes, followed by another spin at 2000 x *g*, 4°C, for 5 minutes. Circulating L-EVs were isolated from platelet-poor plasma by differential centrifugation. Three milliliters of plasma were diluted in the 0.2 µm-filtered PBS to the total volume of 16 mL and centrifuged at 10,000 *g* for 30 minutes (k-factor 2547.2). Resulting crude LO pellets were resuspended in 200 µL of 0.2 µm-filtered PBS and purified from contaminating non-EV proteins by iodixanol density gradient, as described above.

### Whole cell protein lysates

Whole cell lysate (WCL) was obtained after growing cells in serum-free medium for 24 hours and collection of conditioned cell media. Cell monolayers were scraped and washed in chilled PBS ×3 times. Cells were lysed in DTT-free 4% SDS/Tris-HCl lysis buffer^46^.

### Immunoblotting

Immunoblotting analysis was performed as previously described^47^. Protein concentration was measured using Pierce™ 660nm Protein Assay (#22660, ThermoFisher Scientific). The following primary antibodies were used: HSPA5 (#3177, 1:1,000 dilution) and Cytochrome c (#4280, 1:1000 dilution) from Cell Signaling; TSG101 (sc-7964, 1:1,000 dilution), CD9 (sc-13118, 1:1,000 dilution), and Cav-1 (sc-894, 1:10,000 dilution) from Santa Cruz; KRT18 (ab93741, 1:10,000 dilution) and CD81 (ab79559, 1:10,000 dilution) from Abcam.

### Tunable Resistive Pulse Sensing (TRPS) measurements

EV concentrations and size distributions were measured using the qNanoGold instrument (iZON Science, New Zealand). Freshly purified EVs were diluted 1:40 in 0.2-µm filtered PBS. L-EVs (2.8K and 10K) were analyzed using an NP 2000 nanopore (resolution window 0.9–5.7 µm) and S-EVs (100K) were analyzed using an NP250 nanopore (resolution window 110–630 nm). Membranes were stretched at 47 mm and voltage set either at 0.04 V for L-EVs or 0.5 V for S-EVs in order to achieve a stable current baseline of about 120 nA. Particle size and concentrations were calibrated using Izon calibration particles (1:1,000 diluted CPC2000 for L-EVs and 1:100 diluted CPC200 for S-EVs) and a minimum of 500 events were recorded for each sample with a positive pressure of 5 mbar.

### Electrical Sensing Zone (ESZ) measurements

EV concentration and size distribution was also orthogonally measured using the Beckman Coulter® Z Series instrument. Briefly, EV preparations were diluted 1:1000 in ISOTON® II Diluent (Beckman Coulter). The measurements were performed using the following parameters: lower detection limit – 1 μm, upper detection limit – 40 μm, gain – 512, current – 2.828 A, preamp gain – 179.2.

### Transmission electron microscopy (TEM)

EV pellets were analyzed by TEM as described previously^48^. Briefly, EVs fixed with 4% paraformaldehyde were postfixed in 0.5% osmium tetroxide (OsO^4^, Taab, Aldermaston, Berks, UK). The pellets were dehydrated in graded ethanol including block staining with 1% uranyl-acetate in 50% ethanol for 30 minutes and embedded in Taab 812 (Taab). An overnight polymerization of samples at 60 °C was followed by sectioning, and the ultrathin sections were analyzed using a Hitachi 7100 electron microscope (Hitachi, Chiyoda City, Tokyo, Japan) equipped by Veleta, a 2000 × 2000 MegaPixel side-mounted TEM CCD camera (Olympus, Shinjuku City, Tokyo, Japan).

### Label-free proteomic profiling and analysis of cell line-derived EVs

Label-free proteomic comparison of EV samples in biological triplicates was conducted essentially as described^49^. For each replicate, 4 μg EV proteins were digested into tryptic peptides by filter-aided sample preparation^50^. About 1 μg tryptic peptides in 10 μL solution was loaded onto a 2 cm trap column (Thermo Scientific) and then separated on a 50 cm EASY-Spray analytical column (Thermo Scientific) heated to 55°C, using a 3 hour gradient at the flow rate of 250 nL/minute. The resolved tryptic peptides were ionized by an EASY-Spray ion source (Thermo Scientific), and mass spectra (MS1 and MS2) were acquired in a data-dependent manner in an Orbitrap Fusion Lumos mass spectrometer (Thermo Scientific). EASY-IC was used for internal mass calibration. MS1 scans were acquired in 240,000 resolution at m/z of 400 Th, with an ion packet setting of 4 × 10^5^ for automatic gain control and a maximum injection time of 50 ms. Most intense peptide ions with charge state of 2-7 were automatically selected for MS2 fragmentation by higher energy collisional dissociation, using 32% normalized collision energy. MS2 spectra were acquired in the ion trap, using rapid ion trap scan at 1 × 10^4^ automatic gain control and 35 ms maximum injection time. To minimize redundant MS2 acquisition, dynamic exclusion was enabled (repeat count=1, exclusion during=60 s, repeat during=60 s).

The acquired MS data (27 RAW files) were searched against the Uniprot_Human database using MaxQuant (v1.5.5.1). The searching parameters were set as follows: trypsin/P as the protease; carbamidomethyl (C) as variable modification; oxidation (M), acetyl (protein N-term), and deamidation (NQ) as variable modifications; minimal peptide length as 7; up to two missed cleavages; mass tolerance for MS1 was 4.5 ppm and for MS2 was 0.5 Da; identification of second peptides enabled; label free quantification enabled, with match-between-runs within 0.7 minutes. A standard false discovery rate of 0.01 was used to filter peptide-spectrum matches, peptide identifications, and protein identifications.

#### Analysis

In order to standardize the proteomic data, normalization was performed on a per-sample basis.

#### Normalization

Normalization of protein was performed on a per-sample basis, calculated by eq. 1.

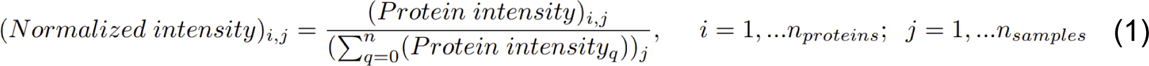

Fraction of proteins in each rank across the nine samples was calculated by eq. 2.

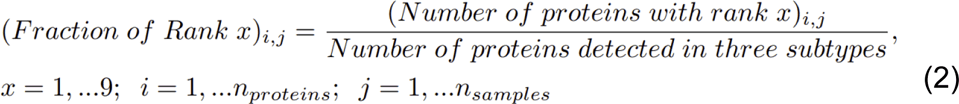

Proteins only detected in one of the triplicates were excluded from statistical analysis. A linear regression model was fit using the R limma package (version 3.42.2). Test statistics from F-test and t-test were moderated by the eBayes function, which applies empirical Bayes method to shrink the protein-wise sample variances towards a common value, thereby augmenting the degrees of freedom for the individual variances. For proteins detected in all three EV subtypes, a multiple linear regression model was fit:

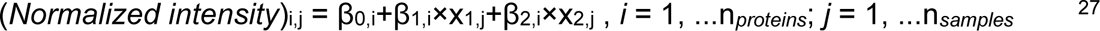

If sample_j_ was in the 2.8K EV subtype, then x_1,j_ = 0, x_2,j_ = 0

if sample_j_ was in the 10K EV subtype, then x_1,j_ = 1, x_2,j_ = 0

if sample_j_ was in the 100K EV subtype, then x_1,j_ = 0, x_2,j_ = 1

For proteins detected in only two EV subtypes (*e.g.* 2.8K pellet and 10K pellet), a linear regression model was fit:

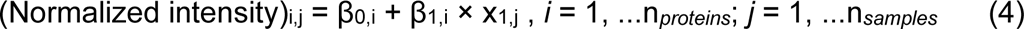

If sample_j_ was in the 2.8K EV subtype, then x_1,j_ = 0

if sample_j_ was in the 10K EV subtype, then x_1,j_ = 1

False discovery rate (FDR) was used to adjust p-values in multiple hypothesis testing. For proteins detected in three EV subtypes, p-values of t-test were adjusted together for all three pairwise comparisons (2.8K *vs.* 10K, 10K *vs.* 100K and 2.8K *vs.* 100K).

All statistical analyses were performed in the programming language R (version 3.6.1). Data was visualized using the BPG library (v5 3.4) ^28^.

#### Statistical Analysis

Statistical analysis was performed based on normalized intensities. Proteins detected in exactly one of the triplicates were excluded from statistical analysis. For proteins detected in all three EV subtypes, a multiple linear regression model was fit to evaluate differential normalized intensity. An F-test was used to test if the mean of normalized intensity was equal across three EV subtypes. A T-test with Welch’s adjustment for heteroscedasticity was then applied to test if the mean of the normalized intensities were equal between every pair of EV subtypes. Threshold of FDR-adjusted p-value < 0.05 from F-tests, FDR-adjusted p-value < 0.01 from t-tests and |log_2_(Fold change)| > 2 were used in identifying differentially abundant proteins.

For proteins detected only in two EV subtypes, a linear regression model was fit and a t-test with Welch’s adjustment for heteroscedasticity was performed to assess the difference in normalized intensity between the two EV subtypes. Threshold of FDR-adjusted p-value < 0.01 from t-tests and |log_2_(Fold change)| >2 were used to identify differentially abundant proteins. The MS data was deposited to the ProteomeXchange Consortium via the PRIDE partner repository with dataset identifier PXD038518.

### SWATH-MS instrumentation and analysis of patient-derived L-EVs

Patient-derived L-EV samples were dehydrated in a speed vacuum and resuspended in 25 μL lysis buffer containing 8 M Urea, 5% SDS and 100 mM DTT in triethylammonium bicarbonate (TEAB). Following brief sonication, samples were alkylated with 10 mM iodoacetamide and processed on Protifi S-TRAP 96-well plates for tryptic digestion according to the manufacturer’s protocol. Eluted peptides were assayed for concentration using the BCA protein assay, dried, and resuspended in 0.1% formic acid in water for injection on LC MS/MS. LC MS/MS analysis was performed on an Orbitrap Exploris 480 (Thermo Scientific, BRE725539) mass spectrometer interfaced with a Nanospray Flex™ ionization source (Thermo Scientific, P/N ES072) coupled to an Ultimate 3000 ultra-high-pressure chromatography system with 0.1% formic acid in water as mobile phase A and 0.1% formic acid in acetonitrile as mobile phase B. Peptides were loaded at 2 µg, based on estimated recoveries from S-Trap processing protocol, and separated on a gradient of 1% B organic phase for 2 minutes, 1-5% B for 0.5 minutes, 5-9% B for 3.5 minutes, 9-27% B for 39 minutes, 27-44% B for 15 minutes, on a C18 column (15 cm length, 300 µm ID, 100 Å pore size, Phenomenex) over the course of 60 minutes with a constant flow rate of 9.5 µL/minute. Source parameters were set to a voltage of 3000 kV, capillary temp of 300°C, and a sweep gas of 2 Arb. MS1 resolution was set to 60,000 and AGC was set to “standard” with ion transmission of 100 ms and RF Lens 40%. Mass range of 400-1000 and AGC target value for fragment spectra of 300% was used. Peptide ions were fragmented at a normalized collision energy of 30%. Fragmented ions were detected across 50 non-overlapping data independent acquisition precursor windows of size 12 Da. MS2 Resolution was set to 15,000 with an ion transmission time of 22 ms. All data was acquired in profile mode using positive polarity.

#### Statistical Analysis

All analyses were carried out in the programming language R (v4.2.1). Consensus clustering (max_k = 10; Spearman’s ρ as the similarity metric; pItem = 0.8; pFeature = 1; seed = 2023; reps = 1000; ConsensusClusterPlus v1.58.0) was performed using hierarchical clustering on 30 samples and the z-scored 75% most variable protein groups that were detected in all samples. The samples from patient A2 are outliers and were therefore, excluded from the analysis. Survival analysis was performed using the R package survival (v3.5.0) to identify the time to progression. Cox proportional models were applied to the before-treatment abundance of the 75% most variable protein groups, adjusting for treatment_ADT and treatment_Docetaxel. C index was used to evaluate the goodness of fit for Cox models. P values were adjusted for multiple comparisons using FDR. Visualizations were created using the R package BPG (v7.0.5). MS data was deposited to the ProteomeXchange Consortium *via* the PRIDE^29^ partner repository with the dataset identifier PXD038011.

### Vesicle flow cytometry

EV concentration, size, and surface marker abundance were measured by single vesicle flow cytometry^51,52^ using a commercial kit (vFC™ Assay kit, Cellarcus Biosciences, La Jolla, CA) and flow cytometer (CytoFLEX™, Beckman Coulter) Briefly, samples were stained with the fluorogenic membrane stain vFRed™ and one or more fluorescent antibodies for 1 hour at RT and analyzed using vFRed fluorescence to trigger detection. Controls included buffer only, reagent only, and positive and negative controls for antigen abundance. Spectral compensation was performed using antibody-stained antibody capture beads and validated using single stained controls. Data were analyzed using FCS Express™ 7 (De Novo Software) and included calibration using a vesicle size and fluorescence intensity standards. The analysis included a pre-stain dilution series to determine the optimal initial sample dilution and multiple positive and negative controls, per guidelines of the International Society for Extracellular Vesicles (ISEV)^29^. A detailed description of vFC™ methods and controls, as well as the MIFlowCyt and MIFlowCyt-EV checklists, as requested by the guidelines are provided in Supporting Information.

### Bulk RNA-seq instrumentation and analysis

RNA was isolated from EV samples using Trizol (Thermo Fisher, Waltham, MA, USA) for RNA sequencing. RNA samples were treated with TURBO DNase (ThermoFisher AM2238) followed by Zymo RNA Clean and Concentrator (R1014). RNA concentration was quantified by Ribogreen reagent (ThermoFisher R11490) and by Agilent TapeStation (5067-5579). Between 2.5-10 ng of RNA were used in sample preparation. cDNA libraries for sequencing were made were made using SMARTer Pico V2 (Takara 634412). Libraries were quantified for paired-end sequencing (101bp x 101bp) on the Illumina NovaSeq 6000 using Kapa Library Quantification Kits (Roche KK4824).

#### FASTQ generation and gene and transcript quantification

FASTQ files were generated using bcl2fastq v2.19.1.403 using default parameters. Reads were trimmed with cutadapt v2.7 according to kit recommendations: -U 3 for SMARTerPicov2. Trimmed FASTQ files were then aligned to GRCh38 primary assembly genome from GENCODE with STAR v2.6.1d with the following parameters: --runMode alignReads --outSAMtype BAM Unsorted --outSAMmode Full --outSAMstrandField intronMotif --outFilterType BySJout --outSAMunmapped Within –outSAMmapqUnique 255 --outFilterMultimapNmax 20 --outFilterMismatchNmax 999 -- outFilterMismatchNoverLmax 0.1 --alignMatesGapMax 1000000 –seedSearchStartLmax 50 --alignIntronMin 20 --alignIntronMax 1000000 --alignSJoverhangMin 18 -- alignSJDBoverhangMin 18 --chimSegmentMin 18 --chimJunctionOverhangMin 18 -- outSJfilterOverhangMin 18 18 18 18 --alignTranscriptsPerReadNmax 50000. Following genome alignment, reads were counted with featureCounts v1.6.3 (part of the subread package) using a non-redundant genome annotation combined from GENCODE 29 and LNCipedia (v5.2) and the following parameters: -p -t exon -g gene_id -s 2. Trimmed fastqs were also quasimapped to the same combined GENCODE 29 and LNCipedia5.2 annotation using salmon quant v0.11.3 to estimate transcripts per million (TPMs) with the following parameters: --libType A --numBootstraps 100 --seqBias --gcBias -dumpEq. Transcript coverage was estimated using bedtools coverage v2.29.0 using default parameters.

#### Analysis

Count data was loaded into R v4.0.3 for analysis. Normalized counts, variance stabilizing transformation, and differential abundance analysis were performed using DESeq2 v1.26.0. Differentially abundant genes with a Benjamini and Hochberg adjusted p < 0.05 and mean-normalized count > 10 were kept for downstream analysis. Heatmaps were created using the ComplexHeatmap (v2.12.1) package in R with Z-scored variance stabilized values as input. UpSet plots were generated using UpSetR (v1.4.0). Pathway analysis was performed on differentially expressed genes using ClusterProfiler (v3.10.1) with Gene Ontology Biological Processes. GSEA was performed using the rank of the Wald test statistic for all genes tested for differential abundance analysis. Data were deposited and approved by GEO (ID GSE214804).

### Single-EV RNA-seq instrumentation and analysis

#### 10x Genomics Chromium library construction and sequencing

L-EVs were isolated from the PC3 cell conditioned medium, purified by iodixanol density gradient, and the number of particles quantified as described above. To enable encapsulation of single L-EVs into individual droplets, the EV sample was diluted to contain 10,000 particles total. A 10x Genomics Chromium machine was used for single-EV capture. cDNA preparations were obtained according to the Single Cell 3’ Protocol recommended by the manufacturer. Silane magnetic beads and solid-phase reversible immobilization (SPRI) beads were used to clean up the GEM reaction mixture, and the barcoded cDNA was then amplified in a PCR step. The P7 and R2 primers were added during the GEM incubation and the P5, and R1 during library construction via end repair, A-tailing, adaptor ligation, and PCR. The final libraries contain the P5 and P7 primers used in Illumina bridge amplification. Sequencing was carried out on an Illumina Novaseq 6000 with paired-end with read lengths of 28 and 91 bp for read 1 and read 2, respectively.

#### Analysis

Data was analyzed using CellRanger (version 6.0.1)^53^ using GRCh38-2020-A provided by 10x Genomics as reference genome. Since default filters are optimized for the detection of cells, we applied custom filters on barcodes and features in the downstream analyses instead, as explained below. Seurat^54^ was used to filter features and barcodes (min.cells=1, min.features=3). Mitochondrial markers are defined by annotation obtained through Biomart^55^ based on human genome hg38. Markers of large oncosomes (LO), small EVs (S-EVs) and LO *vs.* S-EV differentially abundant genes (DEGs) were obtained from bulk RNA sequencing data using the following thresholds:

- LO markers are defined as genes with mean normalized counts = 0 in the 100K fraction and mean normalized counts > 5 in the 10K fraction,

- S-EV markers are defined as genes with mean normalized counts = 0 in the 10K fraction and mean normalized counts > 5 in the 100K fraction,

- LO *vs.* S-EV DEGs were calculated as explained in Bulk RNA sequencing paragraph: briefly, differential abundance analysis was performed using DESeq2 v1.26.0. Differentially abundant genes with a Benjamini and Hochberg adjusted p < 0.05 and mean normalized counts > 10 are retained. All the mitochondrial markers are excluded. Co-occurrence of LO and S-EV marker reads with mitochondrial reads was performed using Fisher’s Test on raw counts, testing different thresholds of detection for mitochondrial reads (from 1 to 20 reads). Data were deposited and approved by GEO (ID GSE231846).

For dimensionality reduction and clustering analysis we proceeded as follows: we filtered out all barcodes with less than 50 counts (n=9343, we introduce this filter in order to remove signal putatively originating from small nanoparticles or ambient RNA and keep signal from LO: this threshold is conservative but does not take into account the presence of doublets since, to our knowledge, there are no methods for excluding aggregates in single EVs sequencing data), then data was normalized and scaled using the NormalizeData (normalization.method = “LogNormalize”) and Scale functions in Seurat. FindVariableFeatures was used to select most informative features (selection.method = “vst”, nfeatures = 1000), then RunPCA was run using the selected features. Following FindNeighbors (reduction = “pca”, dims = 1:10) and FindClusters (resolution = 0.002, algorithm = 2 and resolution = 0.2, algorithm = 2 for analysis of all drops with reads >=50 and drops positive for LO markers, respectively) were run to find clusters. Finally, UMAP was generated using the RunUMAP function (dims = 1:10, method=“umap-learn”, n.neighbors=30, and min.dist=0.5 or min.dist=1 for analysis of all drops with reads >=50 and drops positive for LO markers, respectively). Markers of each cluster were found using FindAllMarkers (only.pos = TRUE, min.pct = 0.25, thresh.use = 0.25). Visualization was obtained using custom scripts and the ggplot2 and ggforce packages. Statistical analysis was performed using R v4.0.2.

## Supplemental figures and figure legends

**Figure S1.**
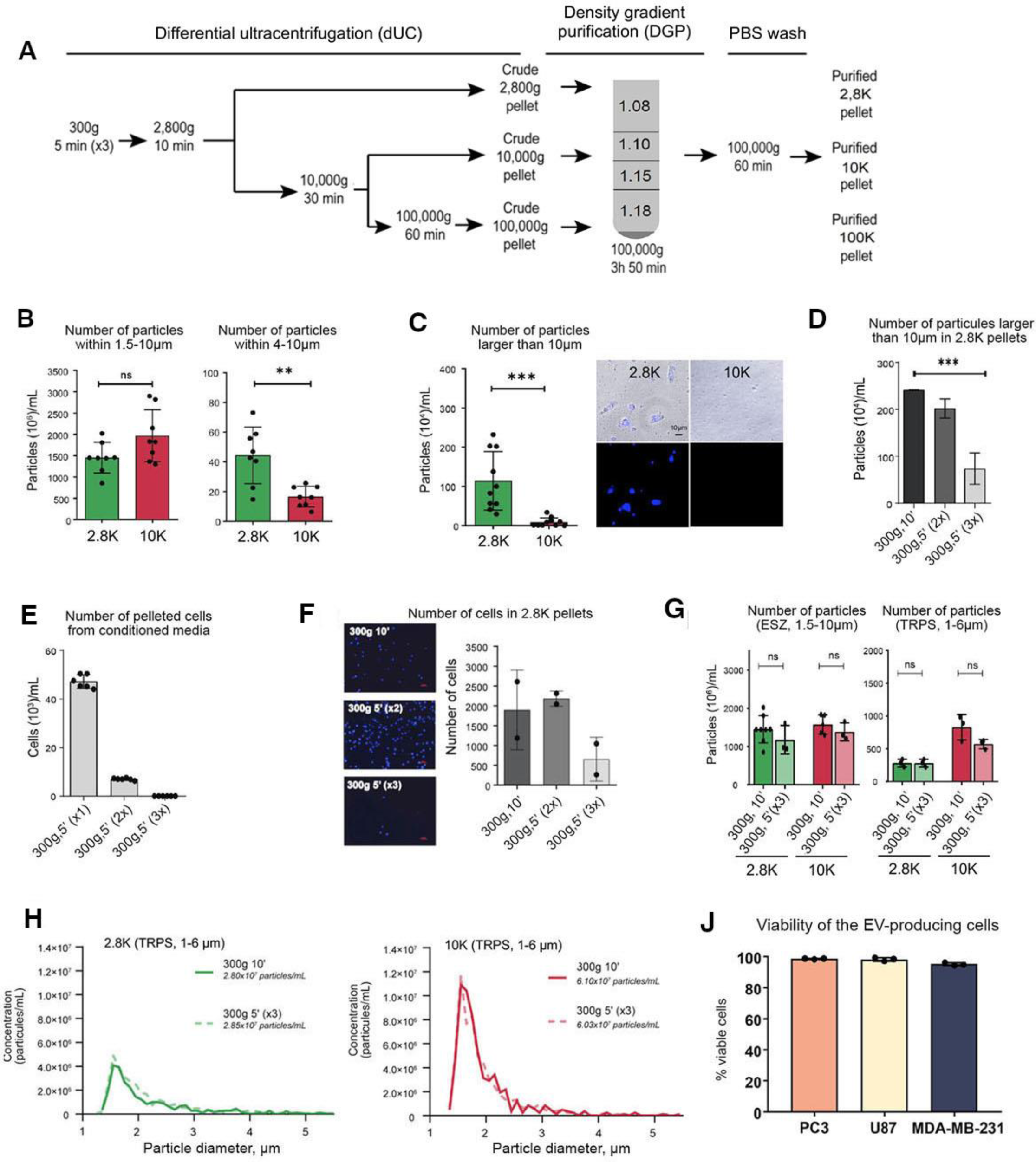
Differential centrifugation protocol optimization to exclude cells from the 2.8K fraction. (**A**) Workflow for purification of three EV populations by differential ultracentrifugation (dUC) at different speeds (2,800g/2.8K, 10,000g/10K, 100,000g/100K) followed by density gradient purification (DGP). (**B**) The electrical sensing zone (ESZ) quantitation of particles within 1.5-10 μm (left) and 4-10 μm^17^ in the 2.8K and 10K pellets obtained from PC3 cell conditioned media after depleting cells at 300g, 10 min. (**C**) (Left) ESZ quantitation of the particles >10 μm in the 2.8K and 10K pellets. Hoechst staining of the 2.8K and 10K pellets cultured for 24 hours showing the presence of some cells in the 2.8K pellet. Scale bar: 10 µm. (**D**) ESZ quantification of the particles >10 µm in 2.8K pellets following different combinations of low-speed centrifugation spins to deplete cells. (**E**) Bar plot showing the number of cells pelleted at sequential centrifugations spins at 300g, 5min, in 50 mL of PC3 conditioned media. (**F**) (Left) Hoechst staining and ^16^ quantification of the cell numbers in the 2.8K pellets following different combinations of low-speed centrifugation spins to deplete cells. Scale bar: 50 µm. (**G**) Cell depletion at 3 x (300g, 5 min) does not affect the recovery of large particles in 2.8K and 10K pellets quantified by ESZ (left) and tunable resistive pulse sensing (TRPS, right). (**H**) TRPS quantification and particle size distribution of the 2.8K and 10K PC3 EVs before and after implementing an optimized protocol for cell depletion demonstrates that the optimized protocol does not alter the recovery of large EVs. (**J**) Average cell numbers and viability in a 150 cm diameter culture dish after 24 hours of serum starvation, prior to collection of the conditioned media for EV isolation.

**Figure S2.**
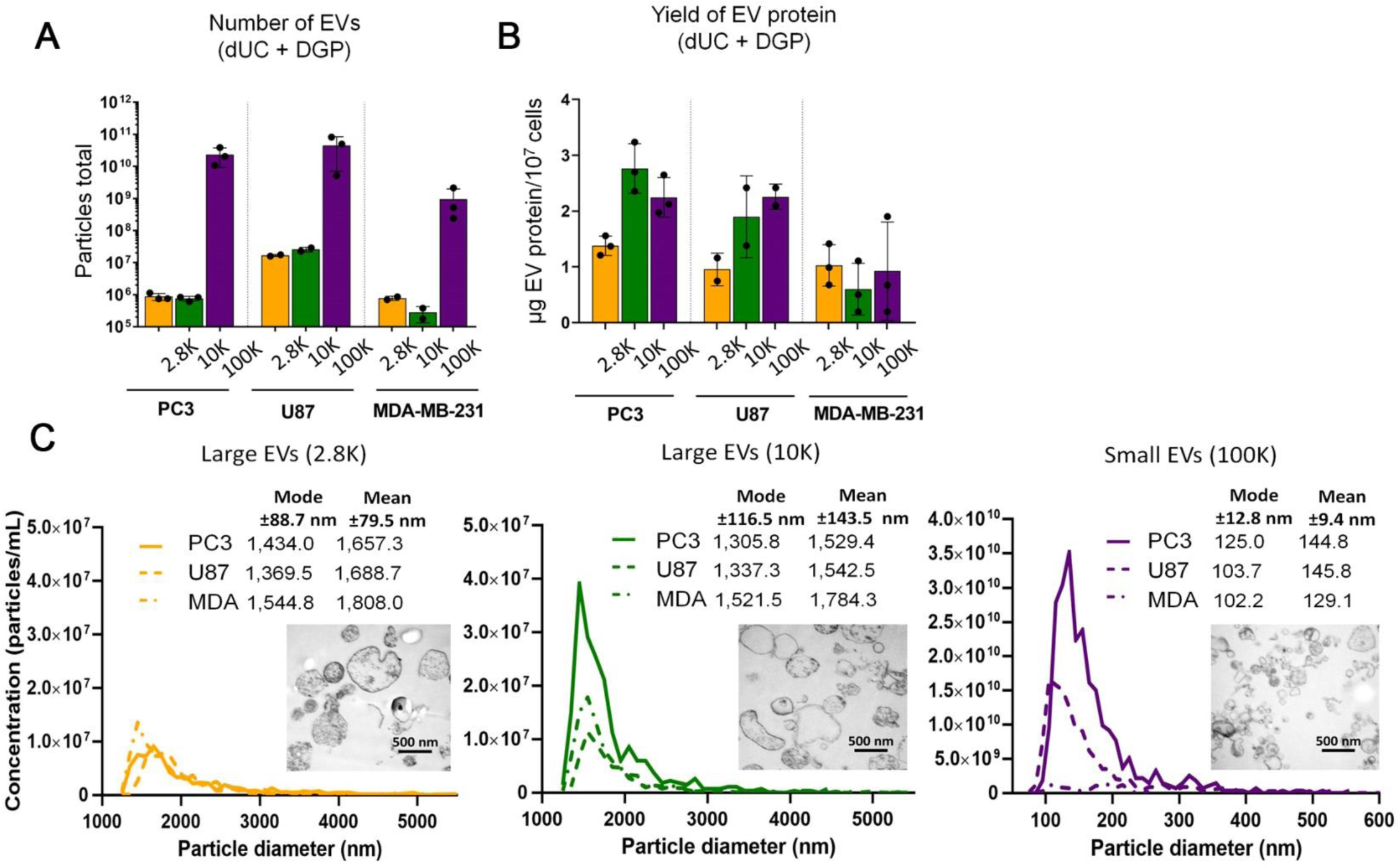
Low-speed centrifugation enriches for a population of L-EVs containing LO. (**A**) Quantitation of the particles by Tunable Resistive Pulse Sensing (TRPS) and EV protein amount (**B**) in the indicated fractions from the indicated cell lines (see methods for details). (**C**) TRPS size distributions of the three EV fractions from the cancer cell lines. Inserts show representative transmission electron microscopy (TEM) images of EVs in each fraction. See also Figure S1.

**Figure S3.**
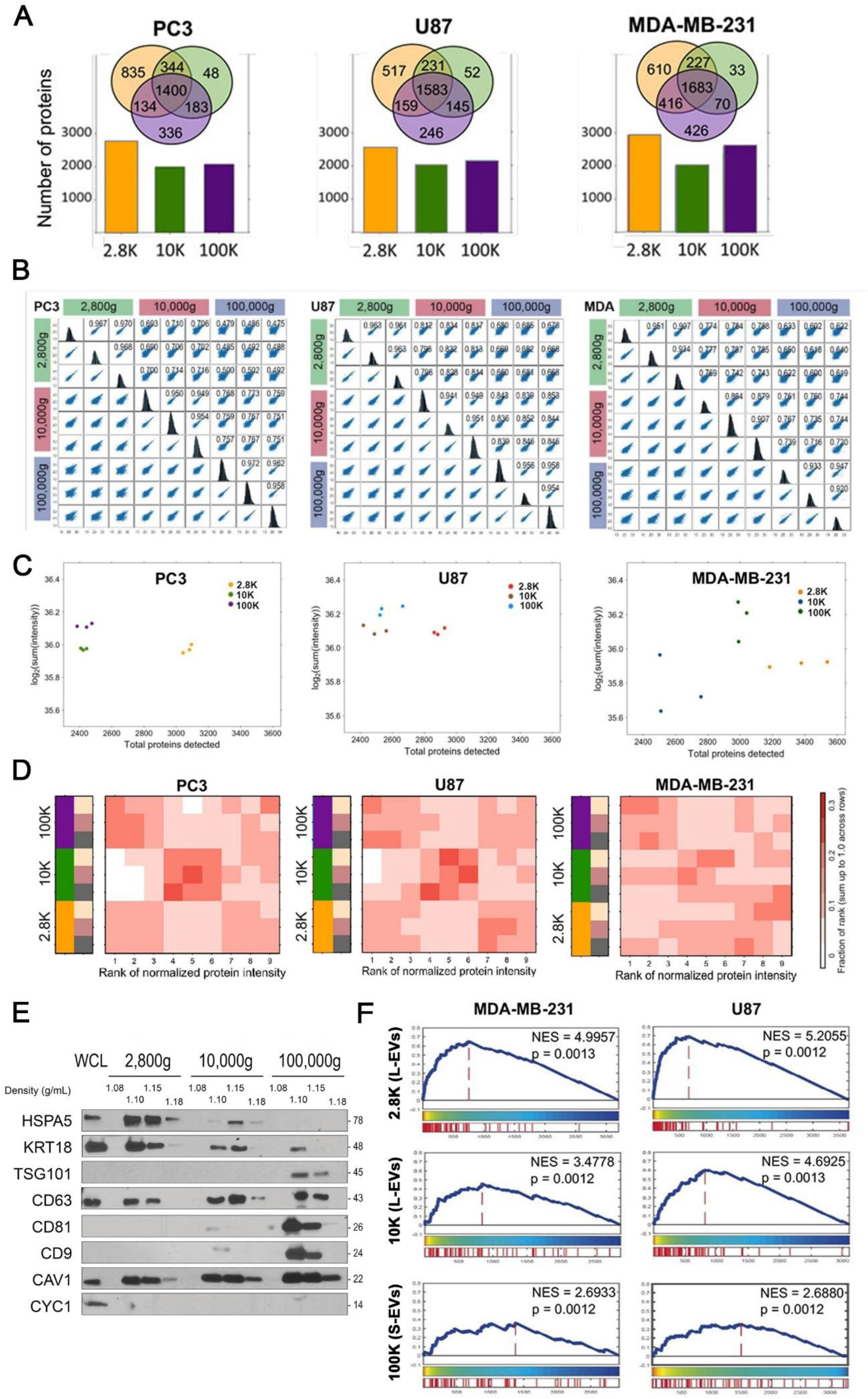
Related to Figure 1. 10K EVs have a distinct protein intensity pattern compared to the 2.8K and 100K EVs. **(A)** Venn diagrams showing the extent of overlapping proteins in each EV fraction (top) from the indicated cells lines. The bar plots summarize the total number of proteins in each fraction (**B**) Spearman’s correlation plots and coefficients of the normalized LFQ signal among technical replicates in the indicated EV populations in PC3, U87, and MDA-MB-231 cells lines (p-value<0.001), and frequency histograms of the normalized LFQ signal for each technical replicate. (**C**) Association between the sum of protein intensity and number of detected proteins in the PC3, U87, and MDA-MB-231 cells lines. (**D**) Rank plots of normalized protein intensity showing a fraction of proteins in each rank across 9 samples for proteins detected in all three EV subtypes in PC3, U87, and MDA-MB-231 cells lines. Each row represents a sample, and each column is the fraction of proteins in each rank. (**E**) The three EV fractions and the whole cell lysate (WCL) from PC3 cells were blotted with the indicated antibodies. (**F**) The gene set enrichment analysis (GSEA) of the proteins differentially enriched in large oncosomes^6^ (FC>1 FDR<0.05) across different EV populations from breast cancer and glioma cell lines.

**Figure S4.**
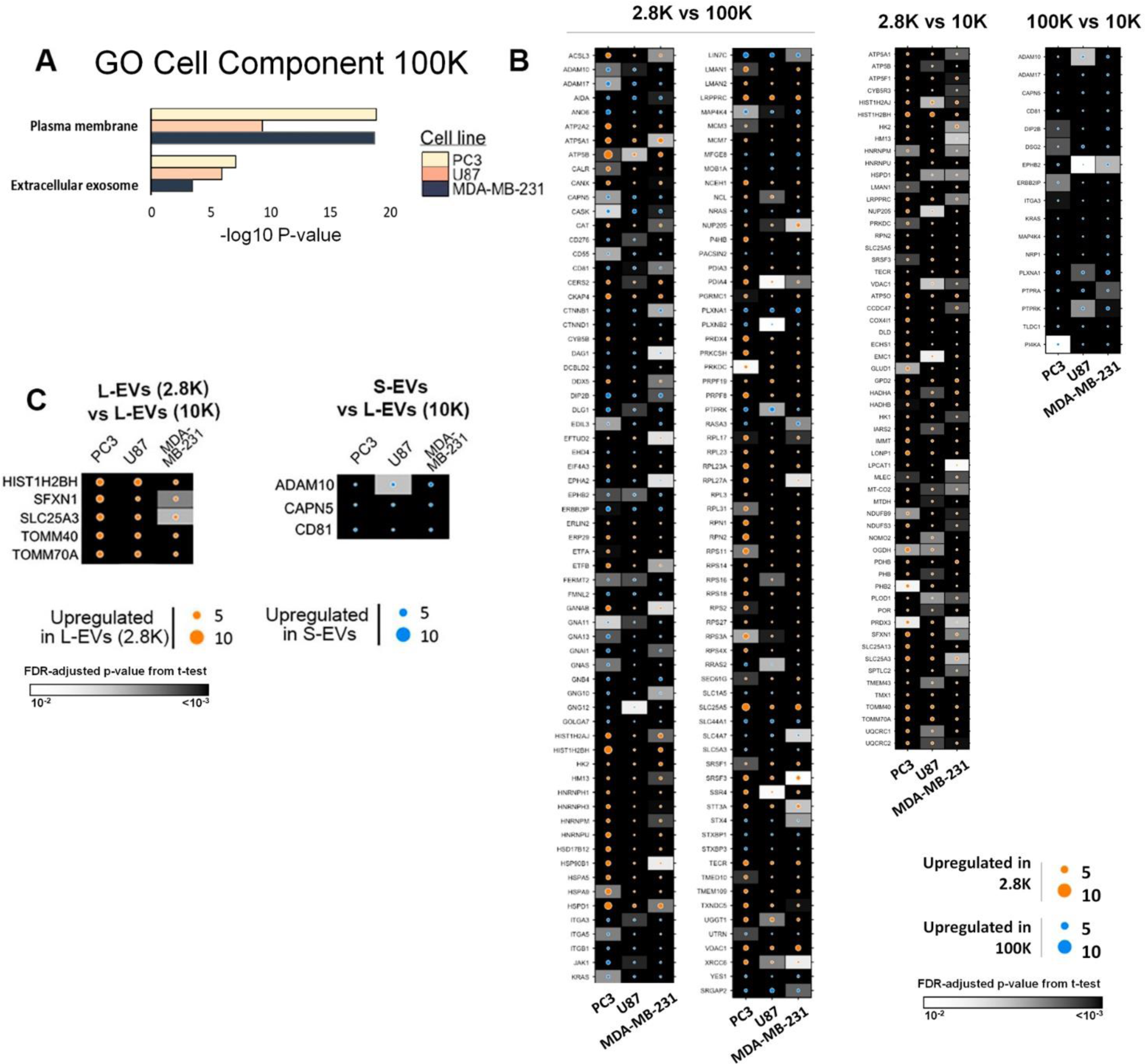
Related to Figure 2. The 2.8K and 100K contain the two most distinct EV populations on the proteome level. (**A**) Gene ontology (GO) analysis for cell component on the proteins unique to 100K demonstrating the enrichment of the plasma membrane and exosome terms across the three cell lines (PC3, U87, and MDA-MB-231). (**B**) Full lists of differentially abundant proteins between the indicated EV fractions that are common to all three cell lines. (**C**) The most enriched (top 25%, fold change > upper quartile) and most highly expressed (top 25%, normalized expression > upper quartile) differentially abundant proteins in 2.8K *vs.* 10K and 100K *vs.* 10K common to the PC3, U87, and MDA-MB-231 cell lines.

**Figure S5.**
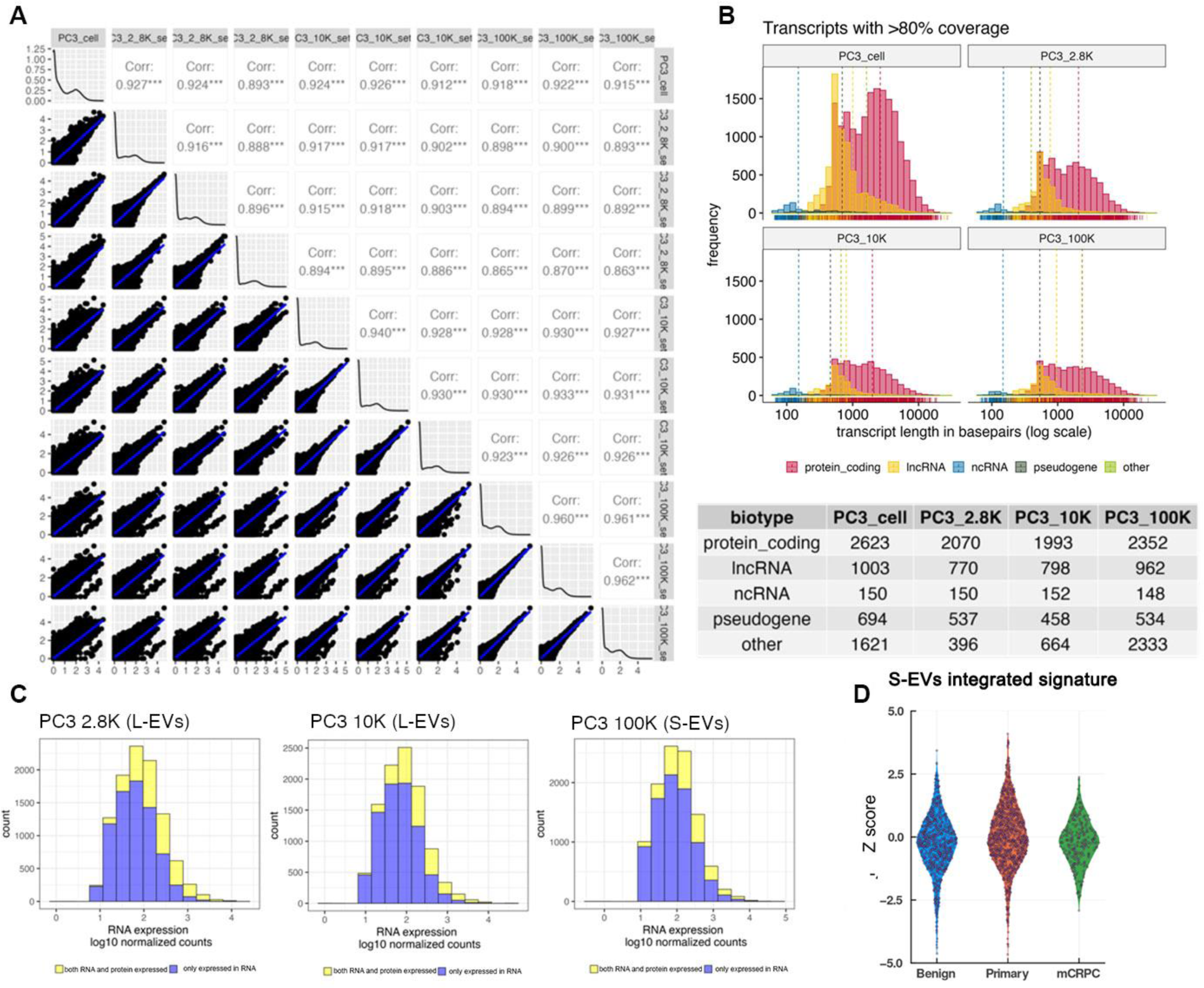
Related to Figures 4 and 6. Transcriptomic analysis reveals a similar RNA composition across the EV fractions. (**A**) Spearman’s rank correlation matrix of transcriptome replicates for the PC3 cell line. (**B**) Histogram of the transcript length of transcripts (TPM > 1) with >80% coverage in different sized EVs from the PC3 cell line. The mean transcript length (bp) is shown in the table below. (**C**) Histogram of genes expressed only in the transcriptome (blue) and both the transcriptome and proteome (yellow) with RNA expression on the X-axis. (**D**) Mitochondrial protein and RNA signature enriched in the PC-derived S-EVs is not significantly upregulated in primary cancer or mCRPC *vs.* benign tissues.

**Figure S6:**
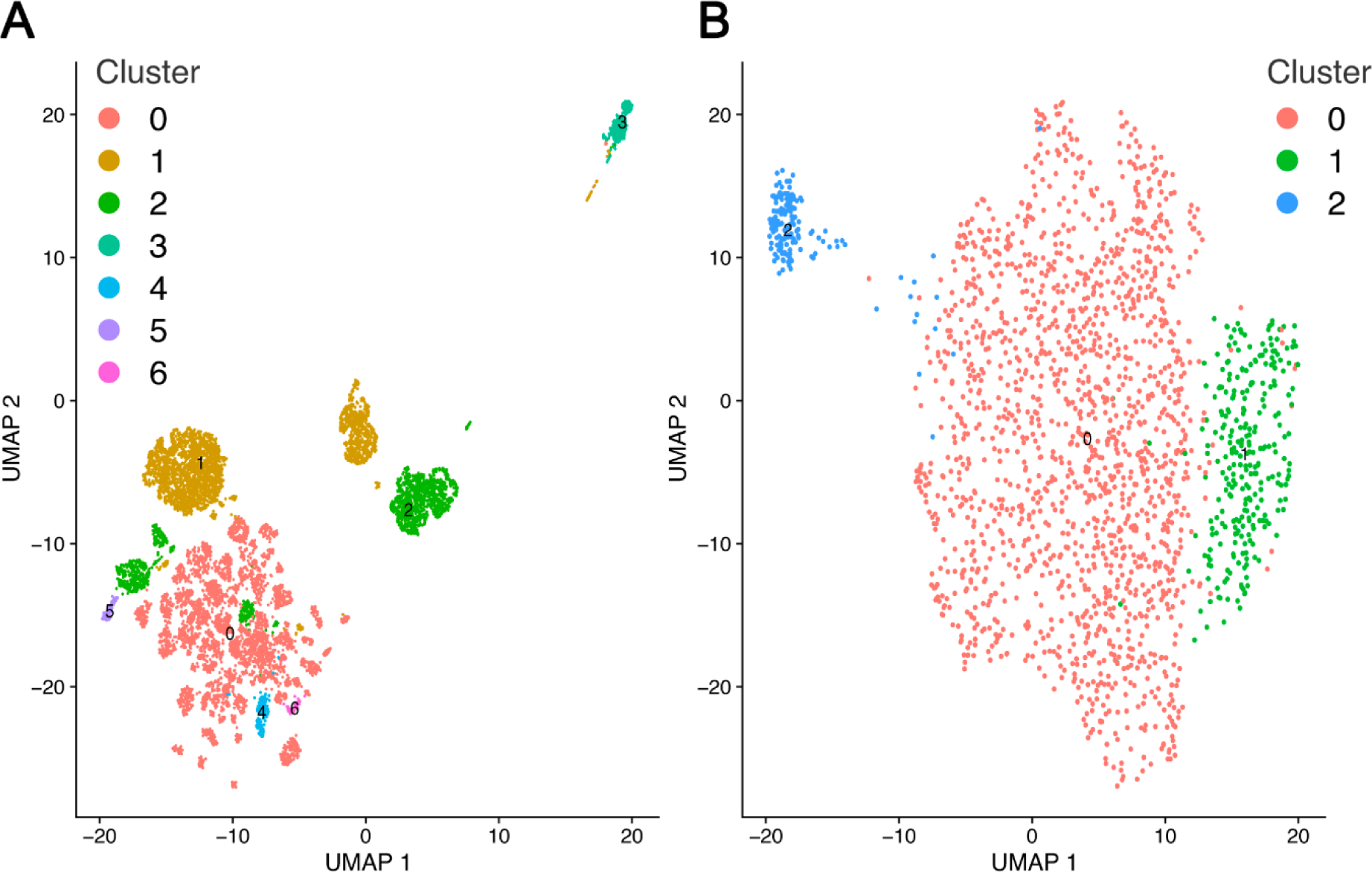
Single-EV RNA-seq analysis of L-EVs. **(A)** Dimensionality reduction of S-EV RNA sequencing data (only drops containing high number (≥50) of reads are shown). Drops are colored by cluster. **(B)** Dimensionality reduction of S-EV RNA sequencing data limited to vesicles positive for LO markers (only vesicles containing high number (≥50) of reads and positive for LO markers are shown). Drops are colored by cluster.

