## Supplemental table 1 for "Extracellular Vesicles heterogeneity through the lens of multiomics"

| Abundant & not detected in non-fractionated human plasma by MS | | | |
| --- | --- | --- | --- |
| Detected only in patient-derived L-EVs | ADCY6 | EPDR1 | TMEM63 |
|  | ATP6V0A1 | F2RL3 | ATMX4 |
|  | CD68 | P2RX1 | TRPC6 |
|  | CD69 | PLXNA4 | TSPAN14 |
|  | CLEC1B | TBXA2R | TSPAN33 |
|  | ENPP4 | TGFB1 |  |
| Detected in patient- and PC3-derived L-EVs | ABCC1 | ITPR1 | SPPL2A |
|  | ABCC4 | ITPR3 | SYPL1 |
|  | ATL1 | NCLN | TMEM30A |
|  | CD151 | PANX1 | TOR1AIP2 |
|  | CD40 | PNPLA6 | TSPAN15 |
|  | CD63 | PTPRA | TSPAN9 |
|  | CRTAP | RPN2 |  |
|  | ERMP1 | SLC44A2 |  |

**Table S1.** Related to Figure 4. Candidate surface proteins in PC-derived L-EVs
